# Unveiling the dimer/monomer propensities of Smad MH1-DNA complexes

**DOI:** 10.1101/833319

**Authors:** Lidia Ruiz, Zuzanna Kaczmarska, Tiago Gomes, Pau Martin-Malpartida, Eric Aragón, Carles Torner, Regina Freier, Błażej Bagiński, Natàlia de Martin Garrido, José A. Márquez, Tiago N. Cordeiro, Radoslaw Pluta, Maria J. Macias

**Author notes:** These authors contributed equally to the work (Lidia Ruiz, Zuzanna Kaczmarska, Tiago Gomes).

## Abstract

R-Smads are effectors of the transforming growth factor β (TGFβ) superfamily and along with Smad4 form trimers to interact with other transcription factors and with DNA. The 5GC-DNA complexes determined here by X-ray crystallography for Smad5 and Smad8 proteins corroborate that all MH1 domains bind SBE and 5GC sites similarly, although Smad2/3/4 MH1 domains bind DNA as monomers whereas Smad1/5/8 form helix-swapped dimers. To examine the relevance of the dimerization phenomenon and to exclude a possible crystallography-induced dimeric state, we studied these MH1 domains in solution. The results show that Smad5/8 domains populate dimers and monomers in equilibrium, whereas their Smad2/3/4 counterparts adopt monomeric conformations. We also found that swapping the loop1 sequence between Smad5 and Smad3 results in the Smad5 chimera-DNA complex crystallizing as a monomer, revealing that the loop1 sequence determines the monomer/dimer propensity of Smad MH1-domains. We propose that the distinct MH1-dimerization status of TGFβ and BMP activated Smads influences the interaction with specific *loci* genome-wide by distinct R-Smad and Smad4 complexes.

**Significance:** TGFβ- and BMP-activated R-Smads were believed to have different preferences with respect to the recognition of DNA motifs and to respond to specific activation inputs. However, recent results indicate that several types of R-Smads can be activated by similar receptors and that all Smads might recognize various DNA motifs. These results pose new questions as to why different types of R-Smads have been conserved for more than 500 million years if they could have a redundant function. They also raise questions as to how different Smad complexes recognize specific clusters of DNA motifs genome-wide.

Here, using structural biology approaches, we elucidate some of the rules that help define the dimers of Smad-DNA complexes and propose how dimers and monomers could influence the composition of Smad complexes, as well as the recognition of specific *cis*-regulatory elements genome-wide.

**Highlights:** R-Smads and Smad4 interact with 5GC and GTCT sites using a conserved binding mode.

Functional differences of TGFβ- and BMP-activated R-Smads are not exclusively related to DNA specificity.

Dimer/monomer propensities are detected in solution and in the absence of DNA.

Loop1 sequence determines the propensity of R-Smads to form monomers or dimers in complexes with DNA.

**Author Contributions:** L.R., Z.K., and T.G. designed and performed most experiments and coordinated collaborations with other authors. L.R. and E.A. cloned, expressed and purified all proteins, L.R., R.F., C.T., N.M., and J.C performed EMSA experiments. L.R., E.A., T.G., T.C., P.M.M., and M.J.M. performed the SAXS and NMR measurements and analyzed the data. P.M.M. analyzed the clustering of DNA motifs in ChIP-Seq data. Z.K. B.B. and R.P. screened crystallization conditions, collected X-ray data and determined the structures. Z.K., R.F. R.P., T.G., J.A.M., and M.J.M. analyzed the structures. All authors contributed ideas to the project. M.J.M. and R.P. supervised the project. M.J.M. wrote the manuscript with contributions from all other authors.

**The authors declare no conflict of interest.**

**Data Deposition:** NMR assignments and chemical shifts have been deposited in the Biological Magnetic Resonance Data Bank, BMRB entry 27548, and the Small-angle scattering data and models have been deposited in SASBDB, entries SASDE32 (Smad5) and SASDE42 (Smad8). Densities and coordinates have been deposited in the Protein Data Bank, entries 6FZS (Smad5), 6FZT (Smad8), 6TBZ (Smad5_3 chimera), 6TCE (Smad5_gly mutant).

**This article contains supporting information.**

## Introduction

The gene responses activated by the TGFβ cytokine family (a term that includes the transforming growth factor β, bone morphogenetic proteins (BMP), Nodal, Activin and other members) play important roles in embryo development, apoptosis, tissue homeostasis, repair, and immunity ^1,2^. These critical roles demand a high level of conservation and fidelity of the TGFβ signaling elements in healthy organisms ^1^.

The main TGFβ signal transduction mechanism is the Smad pathway, with Smad transcription factors being responsible for the transmission of the signals from the membrane receptor into the nucleus ^3^. Receptor-activated Smads (R-Smads) and Smad4 (Co-Smad) are versatile proteins. They all contain a DNA-binding domain (MH1) and a protein-protein interaction region composed of the linker and the MH2 domain ^4,5^. The MH1 and MH2 domains are highly conserved across Smad proteins and along evolution, whereas the linker has a higher sequence variability. R-Smad linkers contain PY motifs and phosphorylatable Ser/Thr residues, which are recognized by cofactors containing pairs of WW domains (Supplementary Figure S1A) ^6,7^. After being phosphorylated at the MH2 domains by TGFβ receptors, activated R-Smads interact with Smad4 and define the canonical hetero-trimeric functional unit. Once in the nucleus, and upon linker phosphorylation, the hetero-trimeric Smad complex is ready to define a new set of interactions with cofactors and with *cis*-regulatory elements, interactions that go on to modulate the outcome of the signaling network ^8-10^.

R-Smad proteins were believed to have different specificities regarding the recognition of DNA motifs and to respond to specific BMP- and TGFβ-activation inputs ^11^. Initial hypotheses suggested that the TGFβ-activated Smads (Smad2/3) and Smad4 showed a preference for the GTCT site (known as the Smad Binding Element, SBE), whereas the BMP-activated Smads (Smad1/5/8) preferred GC-rich sequences. However, the sequence conservation of the MH1 domains (Supplementary Figure S1B) and recent experimental evidence indicate that the separation between DNA binding preferences of R-Smads is subtler than initially thought. For instance, combined TGFβ and BMP receptors influence Smad1/5-driven responses ^12^ and the MH1 domains of Smad3 and Smad4 proteins interact —efficiently and specifically— with a 5GC consensus GGC(GC)|(CG) sequence ^13^. This 5GC consensus is functionally relevant for TGFβ-activated Smads and for Smad4, and it overlaps with the palindromic and compressed 6-BRE site GGCGCC, previously defined as the GC-rich target sequence of BMP-activated Smads ^14^. Crystal structures of Smad2/3 and Smad4 bound to GTCT and 5GC sites, as well as those of Smad1 and Smad5 bound to the GTCT site, have been determined ^13,15-18^. These structures reveal that all R-Smads and Smad4 MH1 domains are able to interact with the SBE, as well as with specific 5GC sites *in vitro* and *in vivo* ^13^. In all these structures, the Smad proteins interact with the 5GC and SBE sites using a distinctive binding mode. Notably, while keeping the same DNA binding characteristics, these crystal structures showed that Smad3 and Smad4 MH1 domains adopt closed conformations ^13,15,18^, whereas SBE-bound Smad1 and Smad5 domains swap the α1 helix between two monomers to form a homo-dimer ^16^. Smad5 bound to the palindromic compressed BRE-GC DNA has also been studied ^17^. Although some structural elements are not well-defined in the model, the protein-DNA interaction is traceable. This binding interface is highly distorted and shows the fewest specific hydrogen bonds between protein and DNA of all Smad complexes determined to date.

In the search for new clues to clarify how BMP-activated Smad proteins interact with GC sites and to decipher the characteristics that define monomers and dimers of MH1 domains, we used X-ray crystallography and other biophysical techniques to study the interaction of Smad5 and Smad8 MH1 domains with non-compressed GC sites. These interactions differed from that described for the BRE-GC DNA. The analysis of these complexes and of the domains in solution allowed us to clarify the molecular basis of the dimer formation, which is dependent on loop1 length and sequence. After swapping the short loop1 sequence of Smad5 for that of Smad3 (slightly longer), we shifted the monomer/dimer equilibrium towards monomeric protein states, while conserving the DNA binding mode. We scanned the motif distribution in ChIP-Seq peaks of Smad-bound regions in regulatory elements of the genome. We observed specific site clustering and motif spacing, depending on whether the regions were BMP- or TGFβ-responsive, as expected for the distinct structural requirements of these Smad complexes. The capacity of MH1 domains to form monomers or dimers can help define the selection of the R-Smad components for a given R-Smad/Smad4 ternary complex, as well as the selection of binding sites in promoters and enhancers.

## Results

### Smad5 and Smad8 complexes with the 5GC site

We first examined how Smad5 and Smad8 recognize 5GC sites and whether the recognition mode is similar to that described for Smad3 and Smad4 ^13,16^ or whether it resembles that reported for the BRE-GC distorted interaction ^17^. To test these hypotheses, we determined the structures of the Smad5 and Smad8 MH1 domains bound to the 5GC GGCGC motif using X-ray crystallography. For the constructs, we used the domain boundaries described in the Smad1/GTCT complex ^16^, which lack the first 10 protein residues since these residues were included in the Smad5/GTCT complex but were disordered ^17^. In both complexes, the GTCT-bound Smad1/5 proteins were oriented as dimers and the interaction with SBE was unaffected by the presence or absence of these additional 10 residues.

Before setting up the crystallization screenings, we studied the protein-DNA interactions by EMSA assays (Supplementary Figure S1C) and observed that the interaction with 5GC motifs was in the same range of concentration as that observed for the SBE sequence. The best diffracting crystals were obtained with a 16bp dsDNA TGCA<u>GGCGC</u>GCCTGCA containing the 5GC sequence (underlined). These crystals diffracted at 2.31 Å and at 2.46 Å resolution for Smad5 and Smad8 MH1 domains, respectively. We solved the Smad5/5GC complex (ASU, space group P2_1_2_1_2_1_) by molecular replacement using a model derived from the Smad1/GTCT complex (PDB: 3KMP) and then used the Smad5 complex to refine that of Smad8 bound to the same DNA. In both complexes, the asymmetric unit contains a dimer of Smad MH1 domains bound to one 5GC site, with the α1 helix being swapped between monomers. The biological unit contains a protein dimer and two DNAs (Smad8 in violet and gray Figure 1A). Crystallization conditions, data collection and statistics are shown in Table 1. The electron densities for the Smad5 and Smad8 proteins and the bound DNA are well defined for the entire complexes (Supplementary Figures S1D,E,F). Smad5 and Smad8 MH1 structures display all the characteristic features of MH1 domains ^5,13^. They are composed of four helices (arranged as a four-helical bundle) and three anti-parallel pairs of short strands (β1-β5, β2-β3, and β4-β6) and the fold is stabilized by the presence of a Zn^2+^ tightly coordinated by three cysteines and one histidine.

**Table 1.** Data collection and refinement statistics.

| GGCGC complex | SMAD5<br>6FZS | SMAD8<br>6FZT | SMAD5_3<br>6TBZ | SMAD5_gly<br>6TCE |
| --- | --- | --- | --- | --- |
| **Data collection** | ESRF-ID30a3 | ESRF-ID30a3 | ALBA-BL13 | ALBA-BL13 |
| Space group | P2<sub>1</sub>2<sub>1</sub>2<sub>1</sub> | P2<sub>1</sub>2<sub>1</sub>2<sub>1</sub> | P3<sub>2</sub> | I4<sub>1</sub>22 |
| Cell dimensions $\square \square \square$<br>a, b, c (Å) | 67.76, 73.37,<br>89.19 | 75.43, 79.51,<br>88.37 | 53.14 53.14<br>83.15 | 106.12 106.12<br>82.40 |
| $\alpha, \beta, \gamma$ (°) | 90, 90, 90 | 90, 90, 90 | 90 90 120 | 90 90 90 |
| Resolution (Å) | 53.96-2.31(2.48-2.31)* | 59.11-2.46<br>(2.63-2.46) | 46.02-1.82<br>(2.08-1.82)* | 53.06-2.92<br>(3.06-2.92) |
| $R_{\text{meas}}$ | 0.130<br>(1.413)<sup>a</sup> | 0.097<br>(1.403)<sup>a</sup> | 0.087<br>(0.885)<sup>a</sup> | 0.049 (3.208)<sup>a</sup> |
| $I/\sigma(I)$ | 11.4 (1.5)<sup>a</sup> | 14.3 (1.3)<sup>a</sup> | 10.9 (1.9)<sup>a</sup> | 21.1 (0.6)<sup>a</sup> |
| $CC_{1/2}$ | 99.8 (52.9)<sup>a</sup> | 99.8 (57.2)<sup>a</sup> | 99.9 (57.8)<sup>a</sup> | 100.0 (31.5)<sup>a</sup> |
| Spherical completeness<br>(%) | 83.0 (22.0)<sup>a</sup> | 79.7 (22.4)<sup>a</sup> | 48.2 (7.3)<sup>a</sup> | 90.1 (34.6)<sup>a</sup> |
| Ellipsoidal completeness<br>(%) | 92.4 (44.6) | 92.0 (54.3) | 88.8 (68.2)<sup>a</sup> | 91.9 (40.6) |
| Redundancy | 9.3 (9.9)<sup>a</sup> | 6.0 (6.3)<sup>a</sup> | 5.0 (3.9)<sup>a</sup> | 7.9 (7.9)<sup>a</sup> |
| **Refinement** |  |  |  |  |
| Resolution (Å) | 29.77-2.31 | 59.11-2.46 | 46.00-1.78 | 53.11-2.92 |
| No. reflections | 16665 | 15793 | 11400 | 4823 |
| $R_{\text{work}} / R_{\text{free}}$ | 0.193/0.241 | 0.193/0.237 | 0.193/0.226 | 0.210/0.253 |
| No. atoms | 2815 | 2852 | 1702 | 1167 |
| Protein | 2010 | 2057 | 988 | 842 |
| DNA | 656 | 656 | 656 | 324 |
| Zinc ions | 2 | 2 | 1 | 1 |
| Water | 143 | 123 | 57 | 0 |
| $B$ factors | | | | |
| Protein | 48.27 | 58.72 | 51.29 | 141.57 |
| DNA | 88.88 | 116.19 | 85.68 | 116.17 |
| Zinc ions | 40.51 | 53.18 | 27.44 | 156.80 |
| Water | 48.06 | 55.94 | 35.78 | NA |
| **R.m.s. deviations** |  |  |  |  |
| Bond lengths (Å) | 0.010 | 0.010 | 0.008 | 0.010 |
| Bond angles (°) | 1.01 | 1.04 | 0.92 | 0.97 |

\* anisotropic correction, (spherical and ellipsoidal)
<sup>a</sup>Values in parentheses are for the highest resolution shell.
Ramachandran outliers: 0% (Smad5, Smad8, Smad5\_3) 1.7% for Smad5\_gly.
Ramachandran preferred/allowed: 98.39/1.61 Smad5 and 96.84/3.16% Smad8, 97.3%/2.7% for Smad5\_3ch, and 88.2%/10.1% for Smad5-gly.

**Figure 1.**
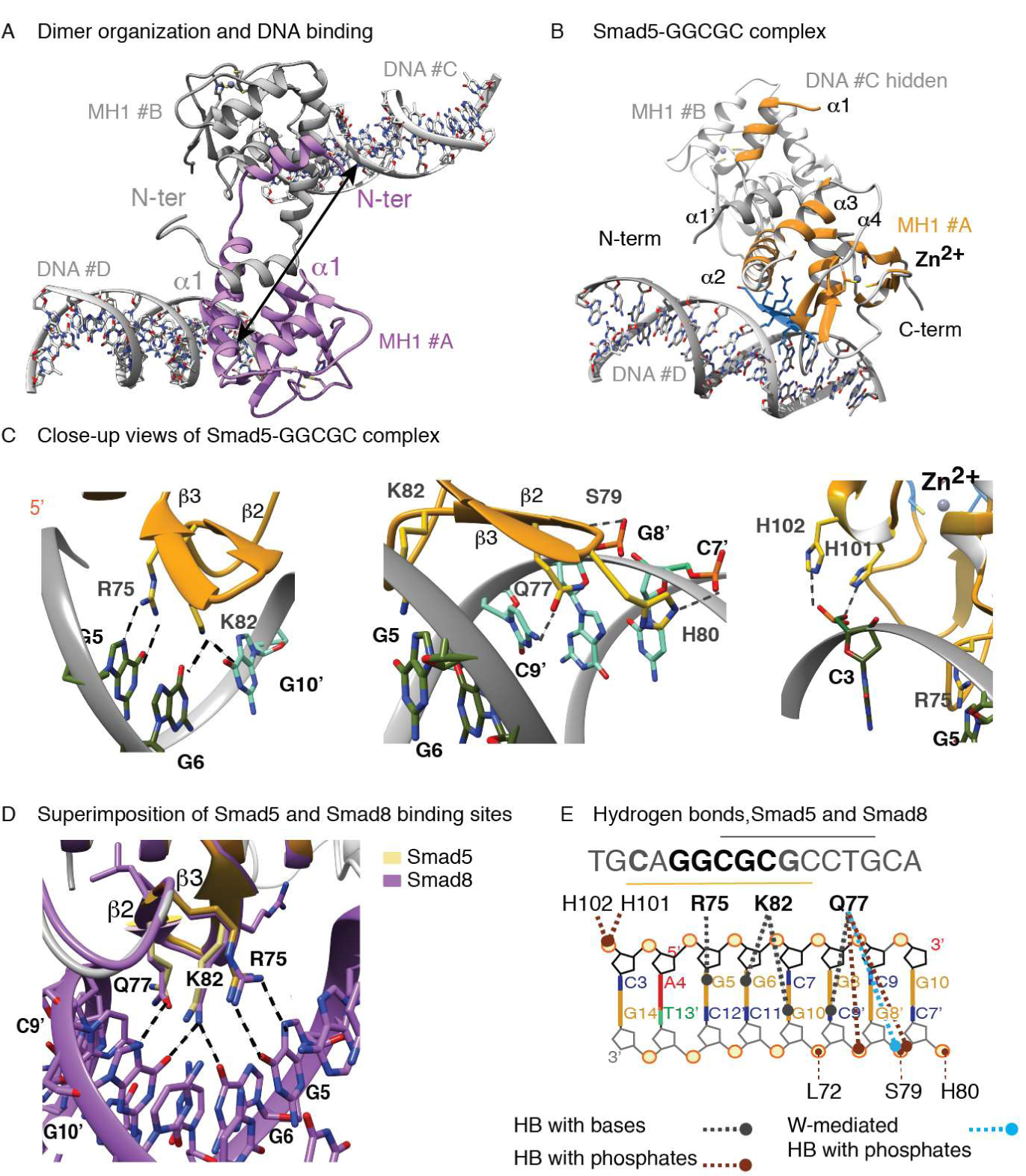
Structures of Smad5 and Smad8 bound to the GGCGC site. A. Ribbon diagram of the homodimeric Smad8 MH1 in complex with the GGCGC DNA motif. Monomers are shown in purple and gray, respectively. The elements of secondary structure and the Zn (gray sphere) binding site are indicated. Each monomer is bound to a different DNA, the distance between DNA binding sites is indicated. B. Ribbon diagram of the Smad5 MH1 domain homodimers in complex with the GGCGC DNA motif (ASU). Monomers are shown in orange and gray respectively. C. Close-up of the Smad5 protein-DNA binding interface. Interacting residues are shown as blue sticks and hydrogen bonds (HBs) as dotted lines. The bases involved in the interaction are labeled. D. Superposition of protein-DNA binding interface for both Smad5 and Smad8. E. Schematic representation of the protein-DNA contacts observed for Smad5 and Smad8 complexes. Grey lines indicate HBs between protein residues and DNA bases whereas brown lines indicate HBs with the DNA phosphates. Blue lines indicate water-mediated HBs with DNA phosphates observed in the Smad5 complex.

The protein-DNA binding region comprises the loop following the β1 strand, and the β2–β3 hairpin (residues 70–83, highlighted in blue, Smad5 complex Figure 1B). This hairpin contains Arg75, Gln77 and Lys82 residues, which are strictly conserved in all MH1 domains. These residues interact directly with the major groove through a network of hydrogen bonds (HBs) with the first four consecutive base pairs of the GGCGCg motif (Figure 1C). The complex is further reinforced by a set of HB interactions between Ser79, Leu72, Gln77, (backbone atoms) and His101 and His102 (side chain) with Gua8’, Gua10’ and Cyt3 bases (Figure 1C, middle and right). There is also a set of nine well-ordered water molecules bound at the interface of the protein-DNA-binding site that contribute to the stability of the complex (Supplementary Table S1). Similar interactions are observed for the Smad8-5GC complex (Figure 1D). When superimposed, the Smad5 and the Smad8 MH1 domains are nearly identical (Cα RMSD of 0.25 Å for 124 aligned residues) and the complexes are very similar to that of Smad1 bound to the GTCT site (3KMP, 123 aligned residues, Cα RMSD of 0.30 Å, Supplementary Table S2). The observed contacts are collected as a cartoon in Figure 1E, showing that one bound MH1 domain covers the 3-CAGGCGC-9 area.

Overall, these results show that homodimers of Smad5 and Smad8 MH1 domains interact with the DNA using a conserved binding mode. This observation corroborates that dimers are a characteristic of these MH1-DNA complexes. MH1 and MH2 domains are connected by a long loop and, given the sequence conservation of Smad1/5/8 at the MH1 dimer interface (Supplementary Figures S1G), we predict that homo- and hetero-dimers of BMP-activated Smads can occur in a cellular context.

### SBE/5GC sites: One binding mode for all Smads

With the exception of the monomer/dimer arrangement of the MH1 domains, the protein-DNA binding interface of the Smad5 and Smad8 complexes is very similar to those of Smad4 and Smad3 bound to SBE and to the same GGCGC motif (Figure 2A) ^13^. The similarity of 5GC is reflected by the conserved pattern of interactions between the protein and the DNA and by the RMSD value of their Cα superimposition (Supplementary Table S2). Even the general DNA topology of the major groove (the principal binding site of all complexes) is conserved between the different bound 5GC DNAs (Supplementary Figure S2), as characterized using Curves ^19^. These complexes also revealed that one MH1 domain is efficiently accommodated on one full DNA major groove, with a clear distinction of minor and major grooves observed in all SBE and 5GC Smad complexes, without introducing protein-DNA structural clashes or distortions. This is in contrast to the arrangement observed in the BRE-GC Smad5 complex ^17^, which shows a conformational-distorted DNA, with both minor and major grooves showing a similar depth and width (Supplementary Figure S2). At first glance, the sequences of BRE-GC (GGCGCC) and 5GC (GGCGCG) appear to be remarkably similar. However, the BRE motif is a 3bp palindrome, and in the crystal two MH1 domains are bound to the BRE-GC site (one MH1 domain is bound to each half of the palindrome) (Figures 2B,C). This effect causes both a huge geometrical perturbation in the B-DNA and a reduction of specific HBs with the protein due to steric hindrance (Figures 2D). Considering that Smad proteins bind to *cis-regulatory* elements containing clusters of motifs ^13^, we believe that the most probable binding mode *in vivo* is that observed in the 5GC and SBE complexes. It seems very unlikely that two MH1 domains would interact with a reduced BRE motif —using half of their protein binding site and causing a high distortion to the DNA structure— if there is the possibility to interact with neighboring sites (as shown in goldenrod in Figure 2B,C) using the full protein binding interface and a perfect accommodation to the DNA.

**Figure 2:**
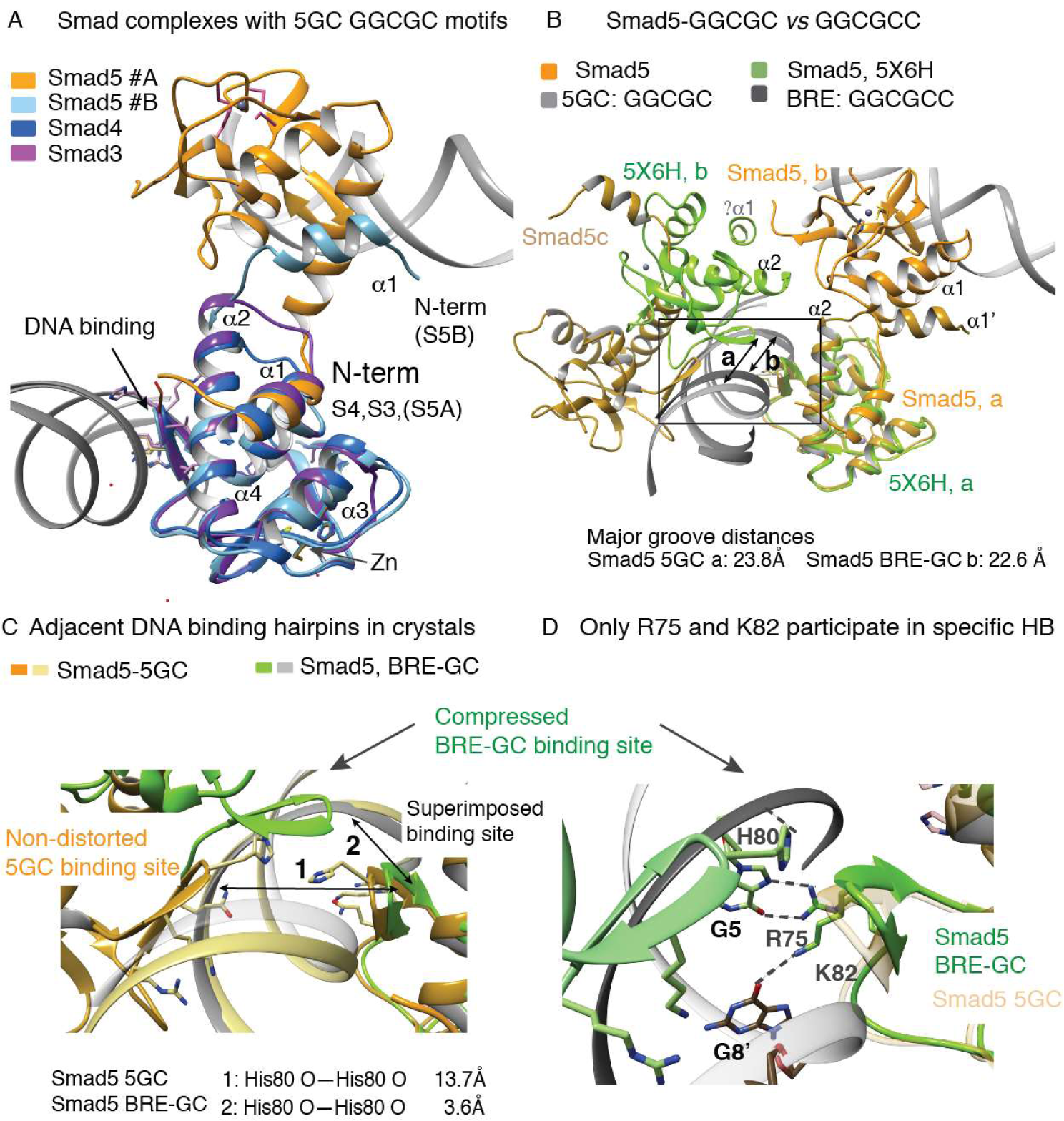
Structural comparison of different Smads bound to DNA. A. Overlay of the complexes of Smad5 MH1 (dimer, gold and light blue) and the monomers corresponding to Smad4 (royal blue) and Smad3 (purple) complexes. All backbones are shown as ribbons. Some secondary structural elements are labeled. N-term of all proteins are indicated. B. Comparison of the Smad5-GGCGC structure (orange) to that of Smad5-BRE (green, PDB:5×6H). Major groove distances are indicated to highlight the DNA compression observed in the Smad5-BRE complex. The first helix of the Smad5-BRE is drawn as described ^16^ although the electron-density map for this region is not well defined. C. Comparison of adjacent DNA binding hairpins in both Smad5 complexes. Distances between the hairpins are indicated. D. Close-up view of the comparison of the DNA binding site. Only the interactions of R75 and K82 with DNA are comparable in both structures (right).

### Smad5/8 MH1 domains display a dimer/monomer equilibrium in solution

We hypothesized that the presence of ensembles of conformations in Smad1/5/8 MH1 domains in solution would facilitate the interchange of structural elements between monomers and dimers in solution resulting in dimers in crystals. This interchange of elements may be promoted by the short length of loop1, which can act as a hinge that directs the helix α1 interchange between Smad1/5/8 monomers. Such loop-length dependent effect has been previously observed as one of the driving forces in formation of domain swapped dimers ^20^. To analyze this hypothesis, we sought to apply nuclear magnetic resonance (NMR) and small-angle X-ray scattering (SAXS). To further characterize the presence of different species, we complemented these studies in solution with data in gas phase provided by ion mobility spectrometry coupled to mass spectrometry (IMS-MS or IM-MS). This technique separates ions, including protein ions, based on their differential mobility through a buffer gas followed by mass spectrometry, therefore allowing separation of complex mixtures, and resolve ions that may be indistinguishable by mass spectrometry alone. Specifically, it allows the separation and differentiation of mixtures of protein complexes since ions of differing shapes show distinct motility—a parameter determined by the characteristic collision cross-section (CCS) of each ion^21^.

We focused the NMR and SAXS parts on Smad5 MH1 domain since both Smad5 and Smad8 have a similar behavior in crystals. We first measured longitudinal, transverse relaxation times (*T1* and *T2*), and heteronuclear ^1^H-^15^N-nuclear Overhauser effect (hetNOE) to characterize the relaxation properties as well as backbone triple resonance experiments for the Smad5 MH1. The analysis of the carbon chemical values allowed us to identify all elements of secondary structure (Figure 3A). Remarkably, the NOE pattern corresponding to the first helix indicates that, in solution and in the absence of DNA, this helix is slightly shorter and more flexible than expected for a fully compact MH1 domain structure. We also observed carbon chemical values close to random coil, as well as lower heteronuclear NOE values for residues located at the α1 helix, in loop1 and at the beginning of α2 helix, thereby confirming the conformational flexibility of these regions (Figure 3A). We also obtained average correlation times (τ_c_) of 12.1 ns and of 13.5 ns at 850 and 600 MHz respectively in contrast to 9.8 ns for the monomeric Smad2, Smad3 and Smad4 MH1 domains previously characterized ^13,22^. We interpreted the large values obtained for Smad5 as an indication of the presence of mixed conformations, including monomers and dimers, in equilibrium facilitated by the flexibility of the α1 helix.

**Figure 3:**
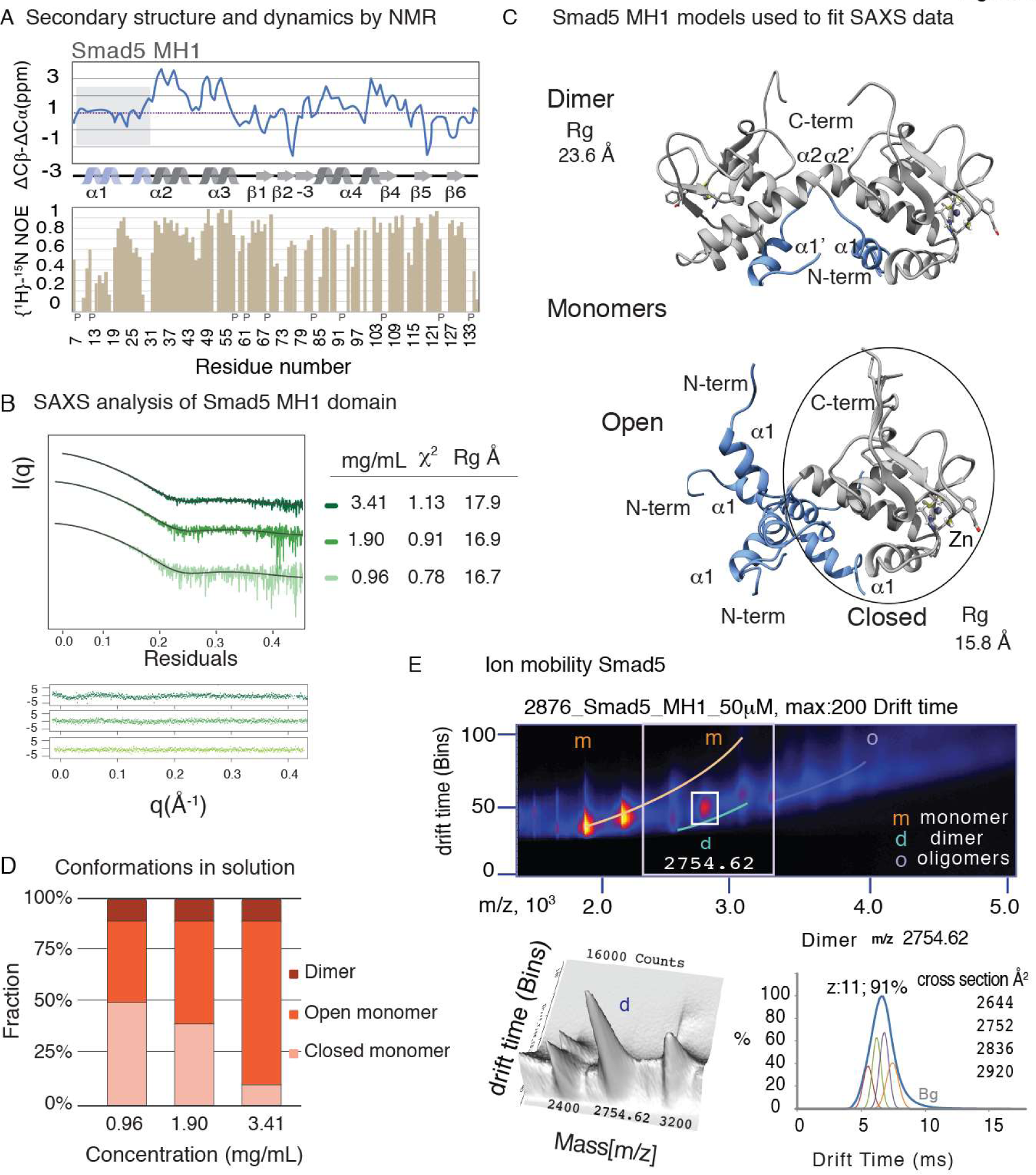
Monomer/dimer equilibrium of Smad5 in solution. A. NMR analysis of the Smad5 MH1 domain. Secondary structure elements based on experimental ^13^Cα-^13^Cβ chemical shift differences with respect to random coil values as well as on characteristic NOEs. Positive and negative values indicate α helices and β sheet structures respectively. The region highlighted in gray represents the values for the first helix, which are close to zero, an indication of variable conformations. Below, hetNOE values at 800 MHz ^1^H Larmor frequency ordered according to the residue number. Proline (P) residues indicated along the sequence. B. Small-angle X-ray scattering (SAXS) analysis of the Smad5 MH1 domains at three protein concentrations. Experimental and graphical output of the best fit shown for each concentration. C. Structural models used for the fitting of the SAXS data. Models were generated from the X-ray structures of Smad5 determined in this work. The closed monomeric structure was generated using the Smad4 structure (PDB: 3QSV) as a template. Trimers were generated with CNS. Since the software did not select MH1 domain trimers for the fitting, they are omitted from the representation. D. Relative abundance of the different conformers in solution. Open and closed monomers and dimers that satisfied the experimental data, indicated as %. E. Plot of the mobility drift time versus m/z for the Smad5 MH1 domain (aqueous solution, 200 mM ammonium acetate buffer) showing the multidimensional data, (instrumental settings are provided in Supplementary Table S3). The mass spectrum from all ions shown in Supplementary Figure S3A. Peaks corresponding to monomers (m) and dimers (d) present in the gas phase are labeled. Below, close-up representation (3D plot) of the conformations corresponding to monomers and dimers that are present at a given m/z value but that are separated by ion mobility IM. IM time distributions for the dimer selected charge state with collision cross-sections used for the fitting (dimer). Calibration curves using standard proteins were used prior to the cross-section analysis.

The presence of several conformations in solution was further analyzed by SAXS (Figure 3B, Supplementary Table S4). Data obtained for Smad5 indicated an interval of radius of gyration between 17.6-19.3 Å, which is comprised between 15.8 Å (expected for a compact monomeric MH1 domain) and 23.6 Å, (the expected value for the fully formed dimer). In order to obtain an accurate fit between the experimental data and the input models, we used oligomeric (trimer and dimer) models together with monomeric (open and closed conformations) models, as suggested by NMR data (Figure 3C). We observed that the relative abundance of the dimer is constant whereas the ratio of open and closed subpopulations within the monomer population varies depending on the sample concentration. The trimer was not selected during the fitting. The best fit was consistent with an equilibrium of monomeric closed and open particles (predominant at 1.9 and 3.4 mg/mL) and swapped dimers (Figure 3D).

Finally, Smad5 dimers and monomers were also studied in the gas phase IM-MS. The analyses of m/z and drift time values revealed that Smad5 can populate monomeric, dimeric, and even higher oligomeric states (trimers and larger), since all these distinct species are resolved at different drift times and can be specifically identified combining the charge state ions of the conformers with their distinct collision cross sections (CCSs). Monomer and dimer forms are more compact, and travel faster than oligomeric ones, which were detected as minority species (Figure 3E, Supplementary Figure S3A-C). The closed and open monomer and dimer conformers have CCS values that correlate well with those of the corresponding structures, as characterized by X-ray diffraction and in solution. Remarkably, dimer/monomer species were detected at all the protein concentrations studied (from 20 up to 150 μM) and under slightly different experimental conditions (Supplementary Table S3). The analysis of the CCS area of selected peaks containing the dimer forms (m/z 2754.6, Figure 3E, and m/z 3029.98, Supplementary Figure S3C) confirmed the presence of several dimeric conformations in Smad5 samples, though monomer and dimer conformations were detected in other peaks (such as m/z 2525.15). Under the same conditions, the most abundant MH1 species of Smad3 and Smad4 were monomeric (Supplementary Figure S3D-E).

All together, these results indicate that, while Smad2/3 and Smad4 in solution, gas phase, and in crystals form monomers, Smad5 (and probably Smad8) populates dimers in equilibrium with monomers, with the dimers being found in crystals of the protein-DNA complexes.

### Replacing the loop1 sequence of Smad5 by that of Smad3 converts Smad5 MH1 domains into monomers

The differences in sequence among Smads are more marked between Smad4 and the R-Smads and to a lesser extent between BMP and TGFβ activated R-Smads themselves. In all Smads, these differences are detected either at the loop connecting the MH1 and MH2 domains or at loops within these two domains ^4^. If we focus on the differences observed at the R-Smads MH1 domains, Smad1/5/8 have four residues in loop1, whereas the same loop has six residues in Smad3 and in Smad4, and sixteen in Smad2 (Supplementary Figure S4A) ^22^. The different lengths of loop1 and also of the α2 helix have an impact on the MH1 structures of the different Smads. When the structures of six Smad MH1 domains were superimposed (Figure 4A), we observed that Smad2/3 long loops can bridge the distance to accommodate the α1 helix packed to the same monomer. These domains have the α2 helix one turn longer than in Smad4 MH1 domain, which is also monomer (Figure 4A, Supplementary Figure S4A). However, it seems that the combination of a long α2 helix (as long as in Smad2/3) with a short loop1 explains the difficulties of the Smad1/5/8 to maintain a fully folded α1 helix and a compact structure (Figure 4B) and instead, in crystals (and in solution of free proteins), the loop1 and the α1 helix protrude away and are swapped between two monomers to form a dimer. In contrast, in the case of Smad3, the turn in loop1 is stabilized by internal hydrogen bond contacts (PDB 5OD6), (Figure 4C) ^16^. In the other monomers, Smad2 has a disordered loop1 due to its long Gly-rich insertion (indicated as an asterisk, Figure 4A, PDB 6H3R) ^22^, whereas in Smad4, the loop is well defined without the presence of internal HBs (PDB 5MEY, Supplementary Figure S4B).

**Figure 4:**
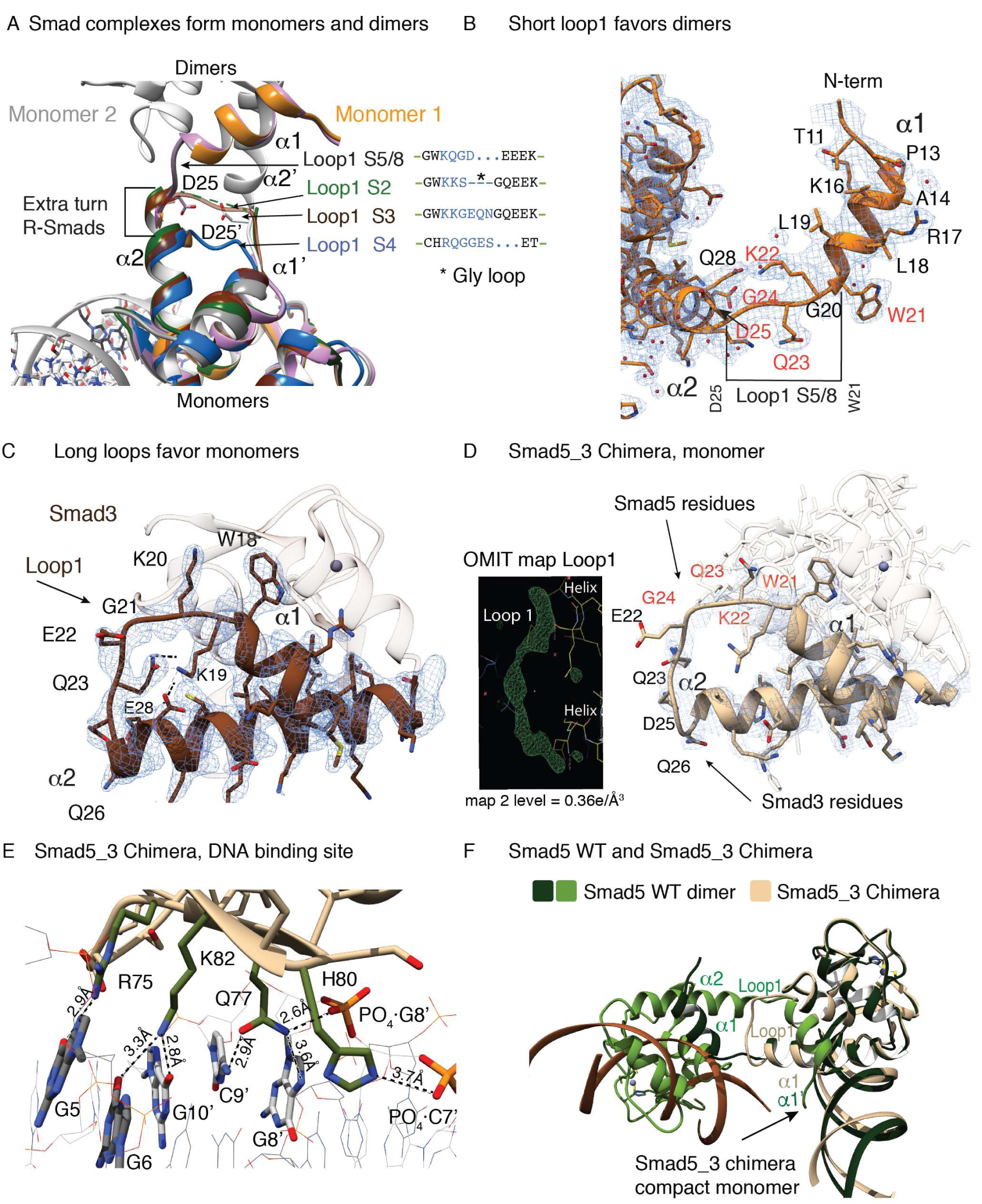
Structural features defining dimer/monomer propensities. A. Close-up of loop1 in the different structures. Differences in loop1 (sequences are indicated) and in the second helix, which is longer in R-Smads than in Smad4, are indicated with arrows and as a bracket, respectively. The positions of D25 for both monomers in Smad5 (the last residue of the α1 helix) are indicated, to highlight the difficulty of BMP Smads to bridge this distance with a short loop. The asterisks in Smad2 denotes the Gly insertion of loop1. B. A section of the electron‐density map for the Smad5 MH1 dimer. To indicate the contacts at the protein dimer interface, some elements of the secondary structure and selected residues are labeled surrounding the α1 helix. Loop1 residues are indicated in red. C. A section of the electron-density map of Smad3 monomer (pdb: 5OD6). The map is shown for helixes 1 and 2 and loop1, (contoured at 1.0σ). Some side chains and the HB stabilizing the turn are labeled. D. A section of the electron‐density map for the Smad5_3 chimera showing the loop1 orientation and the monomer arrangement. Some elements of the secondary structure and selected residues are labeled surrounding the α1 helix. The omit map corresponding to the loop is also shown. E. Close-up view of the Smad5_3 chimera bound to DNA. Residues located at the protein-DNA binding site are shown in light gray. Hydrogen bonds are indicated as dotted lines. F. Superposition of Smad5 dimer and monomer (green and beige). The loop1 in the monomer (chimeric construct) and the dimer interface (Smad5 WT) are labeled. The arrow indicates the compact fold of the chimeric construct.

Similar observations have been reported in the literature, where hinge loop shortening led to dimer formation for proteins that otherwise would adopt monomeric forms. A well-known example corresponds to the Staphylococcal nuclease, where such a loop deletion favors a homodimer association^20-23^. Prompted by these results, we performed the opposite experiment by preparing two Smad5 MH1 mutants with extended loop1. The first construct contained a GGGS insertion in the loop1 (Smad5_gly mutant) and was shown by IM-MS to form mainly monomeric species in the gas phase (Supplementary Figure S3F). Furthermore, Smad5_gly-DNA complex crystallized as a monomer with two MH1 domains bound to two 5GC sites on one DNA molecule (with the same crystal packing as in Smad3 crystals of the PDB_5ODG structure). However, in these crystals the loop1 could not be fully traced in the electron density map due to the flexible nature of the GGGS insertion (Supplementary Figure S4E). For the second mutant (Smad5_3 chimera), we replaced the Smad5 loop1 with that of the Smad3. We obtained well-diffracting crystals of this complex, which allowed us to solve the structure at 1.8 Å (the highest resolution structure so far for MH1-DNA complexes). The complex contains only one MH1 domain bound to the DNA in the asymmetric unit and we observed specific side chain-nucleobase contacts with all five bp of the 5GC Smad-binding site. In this case, the loop1 is well defined in the electron density 2Fo-Fc map (the presence of all residues is further supported by strong positive peaks of the Fo-Fc omit map when deleting loop1 residues before the refinement; Figure 4D, Supplementary Figure S4C,D). Regarding the DNA recognition, the binding mode is conserved with respect to the Smad5 homodimer (and other Smads) with the presence of a new specific HB being formed between Q77 and Gua8’ of the GGCGC motif (Figure 4E). Additionally, Q77 and H80 side chains make HBs with phosphates of Gua8’ and Cyt7’, respectively. The comparison of the chimeric construct and the wild type dimer is shown as Figure 4F.

In summary, these structures confirmed that the propensity to form dimers or monomers is facilitated by the variable length of the loop1 sequence. Whereas the longer loops of Smad2/3/4 (monomers) are more variable in length and sequence, the short Smad1/5/8 loop has been conserved during metazoan evolution, perhaps indicating that the tendency to form dimers is associated with biological function.

### Differential distribution of Smad-binding motifs in BMP- and TGFβ-responsive elements (New section)

As we have observed previously ^13^, Smad-binding sites tend to occur in clusters of three or more Smad-binding motifs. Our hypothesis is that the different Smad4 and R-Smads heterotrimers would interact with Smad binding sites whose separation will depend on the specific R-Smads and on their capacities to form monomers or dimers of MH1 domains.

To assess whether monomer or dimer preferences are reflected in the distribution of motifs within regions bound by Smad complexes in regulatory elements, we sought to analyze ChIP-seq data available in public databases for both BMP and TGFβ-activated Smad proteins. For this analysis, we found two datasets performed in mESC (E14 cell lines) with data obtained for Smad1/5/8 and Smad2/3 proteins ^22,24^. For motif counting, we divided the study into two analyses. We selected peaks found near genes regulated by either TGFβ (Smad3 dataset) or by BMP (Smad1/5 dataset) signaling pathways described in the literature (collected in Supplementary Table S5) or we analyzed all regions with Smad-bound peaks without specific loci localization for each dataset ^25-27^. We normalized all ChIP-seq peaks to be 200 bp, and scanned the set of known Smad-binding motifs (5SBE and 5GC, equal length sequences) to obtain their frequencies in these regulatory regions. When we scanned the set of specific genes, we found that TGFβ-regulated promoters have a higher average count of Smad-binding motifs than the BMP ones (**20** motifs/Kb and **13** motifs/Kb, respectively). When the analysis was extended to all peaks, we observed a similar trend, although the values were slightly smaller than before (**16** motifs/Kb and **9** motifs/Kb respectively), (Figure 5A). We also studied the distribution of the 6-base GC-BRE site for comparison. This motif is (as expected) approximately 4 times less frequent than that of the 5-base 5GC (29% of the 5GC GGCGC motifs are followed by C, which is the BRE motif) and therefore, we could not detect significant enrichment for the Smad1/5 or Smad3 datasets when searching for it. The motif distribution was analysed by the Anderson-Darling normality test, yielding non-normally distributed data as indicated by p-values <0.01, well below the 0.05 threshold value. We also compared both non-normal distributions using the Wilcoxon rank sum test and obtained a p-value =2.2e-16, (much lower than 0.05 threshold), confirming that the differences in motif distribution were statistically significant. We also measured the inter-motif distances for each cluster, calculated with respect to the ChIPSeq-peak centre, and observed that they show no special preference for being localized near the centre or at the boundaries. On average, we observed that the Smad-binding motifs were separated by ∼**46 bp** in BMP-activated regions and by ∼**33 bp** in TGFβ ones (16 bp are equivalent to 50 Å distance), (Figure 5B). These values indicated that BMP responsive genes have clusters of Smad-binding sites more separated than TGFβ ones, in agreement with the observation that Smad3 data have a higher number of motifs/Kb than Smad1/5 data. For comparison, in uniform distributions, the expected values would have been 83 bp for BMP (1000/(13-1 motifs/Kb))=83 bp) and 52 bp for TGFβ ones (1000/(20-1))=52 bp. Again, the data were non-normally distributed (Anderson-Darling test), and the Wilcoxon test showed that the differences in inter-motif distances were significant (p-value under 2.2e-16<<0.05).

**Figure 5:**
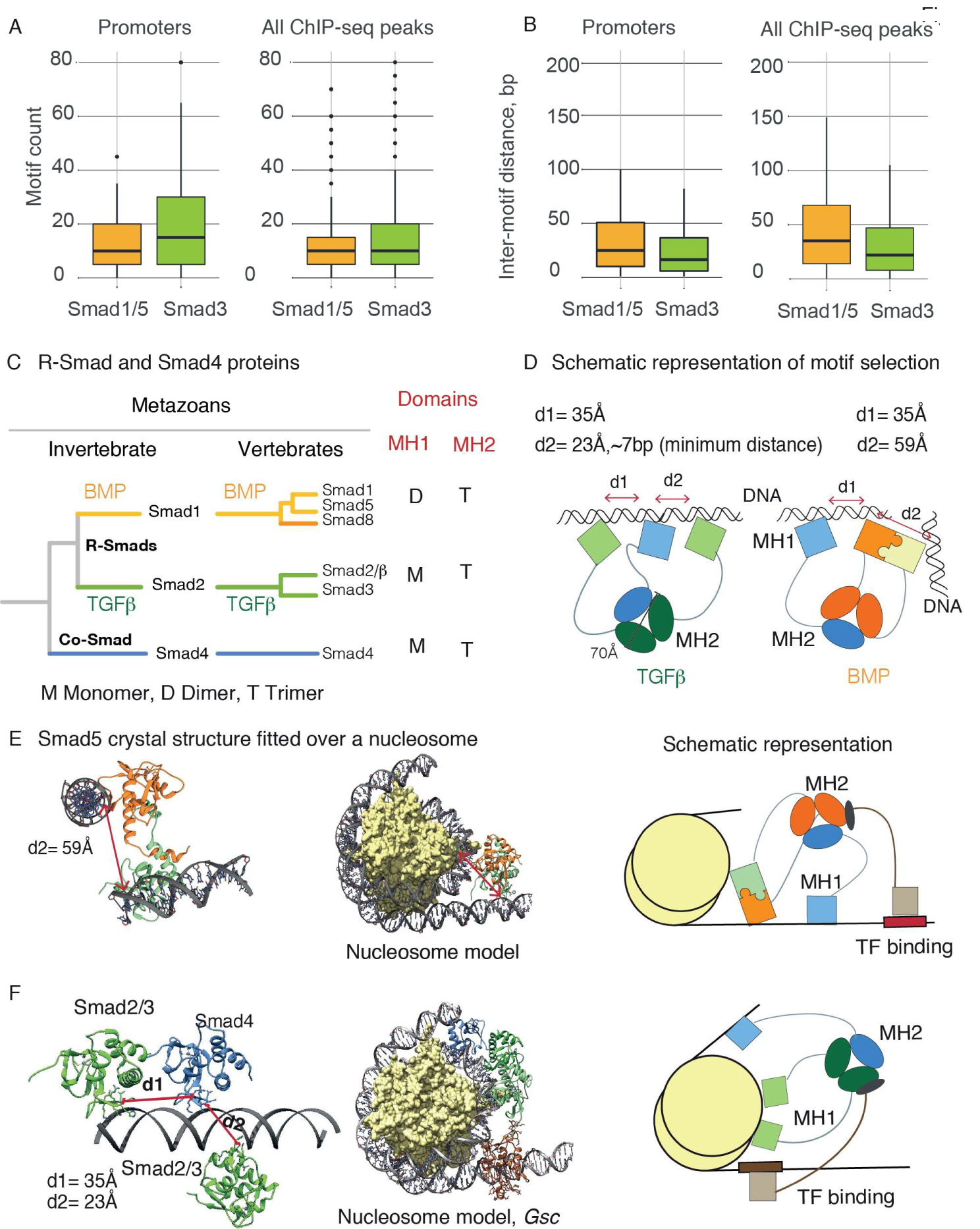
Structure-based models of Smad heterotrimers interacting with DNA clusters. A. Left. Box-plot representing the motif count (SBE+5GC) for Smad1/5 and Smad3 ChIP-Seq datasets, for BMP- (Smad1/5) and TGFβ- (Smad3) regulated promoters. Right. Box-plot of the motif count for all detected peaks for Smad1/5 and Smad3 ChIP-Seqs. The list of promoters is available as Supplementary Table S5. B. Left. Box-plot of the inter-motif distance, in bp, for the motifs (SBE+5GC) for Smad1/5 and Smad3 ChIP-Seqs, for BMP- (Smad5) and TGFβ- (Smad3) regulated promoters. Right. Box-plot of the inter-motif distance for all detected peaks for Smad1/5 and Smad3 ChIP-Seqs C. Schematic representation of the different Smad proteins in metazoans, including the duplications observed in vertebrates, adapted from ^4^. For each R-Smad protein we indicated the propensity of their MH1 domains to fold as monomers and dimers (M, D) and of their MH2 domains= to associate as heterotrimers with Smad4 (T). D. Schematic representation of Smad1/5-Smad4 and Smad2/3-Smad4 heterotrimers binding to SBE/5GC sites modelled based on the Smad binding region present in the *Gsc* promoter. The dimers of MH1 domains would select DNA sites separated by the distance of the two DNA-binding hairpins in the dimer (a distance of approx. 60 Å) and with the Smad4 bound as close as 23 Å (the minimum distance between two consecutive DNA sites that allows the interaction of two MH1 domains without steric clashes between the domains. Furthermore, these dimers can only bind to DNA strands with a certain relative orientation imposed by the dimer structure. Conversely, MH1 Smad2/3 monomeric complexes of Smad2/3-Smad4 heterotrimers are not restricted by the location of their binding sites and can be as close as 7 bp (measured from the center of the motifs). Smad cofactors and other chromatin-interacting proteins are omitted for simplicity. E. Model of Smad5–nucleosome complex. *Left:* Crystal structure of MH1 Smad5 dimer (orange and green) bound to two distant GGCGC sites (6FZS). *Middle* MH1 Smad5 dimer docked manually on a nucleosome (166 bp) structure described in the literature^41^ (6GEJ, in yellow and grey). The two modeled Smad-binding sites are approximately 85 bp away. *Right:* cartoon representation of the full-length Smad-nucleosome-transcription factor complex model. The presence of a transcription factor (TF) cofactor is indicated. F. As in E, but for Smad2/3-Smad4 heterotrimers.

### MH1 domain monomers or dimers, a mechanism for selection of different sites in clusters of SBE/5GC motifs (New section)

The monomer/dimer formation of MH1 domains would also have implications in the selection of several DNA sites recognized simultaneously by the Smad complexes. Heterotrimers with dimeric or monomeric R-Smads might interact with select clusters whose motif distribution fits the separation to best accommodate their MH1 domains. The different distributions might indicate that the BMP-Smad4 heterotrimers bind one Smad4 and one of the two BMP-activated Smads in a given cluster, whereas the second BMP-activated Smad is probably bound at a neighbouring (or distant) one due to spatial constrains dictated by a fixed distance between the DNA-binding hairpins in Smad1/5/8 dimers (Figures 5C,D). For TGFβ-Smad4 heterotrimers, the motif choice is not structurally restricted, and the three Smads can interact within a single cluster of sites (Figure 5D).

To illustrate these hypotheses, we explored the possibility of Smad complexes interacting with nucleosomes, for instance as part of complexes with pioneer factors. We found that the Smad5 MH1 dimer docks well to a linker histone-depleted nucleosome structure (Figure 5E), with the minimal distance between two potential Smad1/5 binding sites being equal to 80-90 bp or larger (hundreds of bp) when the sites are located within nucleosome-depleted DNA or in (nucleosome) linker DNA regions. In contrast, Smad4 complexes with TGFβ-activated Smads (all MH1 domains monomers) can bind both distant (motifs separated by dozens of bp) and/or adjacent motifs within the same cluster, thereby allowing for binding as close as it could occur when binding to the *Gsc* promoter in the presence of FoxH1 (Figure 5F).

In all, dimeric Smad1/5/8 MH1 domains will select DNA sites separated by the distance of the DNA binding sites in the dimer. This property differentiates them from the monomeric MH1 domains, which can interact with motifs whose separation only depends on the steric hindrance between two proteins. Thus, a unique MH1 fold provides two structural/functional solutions (dimers or monomers) to interact with specific promoters and enhancers depending on the cellular context.

## Discussion

In eukaryotes, the association between transcription factors (TFs) to form homo- and hetero-dimers is a common feature employed by many TF families. This capacity of association has implications in the regulation of specific cellular responses, in the stability of the proteins, in the optimal selection of DNA binding sites and in determining overall affinity ^28-30^. Although Smad proteins form trimers via their interactions with MH2 domains, we found that they also follow this rule present in many TF with respect to their association with other Smads through MH1 domain swapping. By doing this multiple association, Smad complexes can fine-tuned their capacity to recognize clusters of binding sites containing motifs separated according to the structural requirements of their MH1 domains (monomers or dimers) ^5,13,31^.

Overall, our results reveal three new hypotheses for the function of Smad complexes *in vivo*. First, the comparison of the Smad5/8 complexes determined here to those previously characterized for Smad2/3 and Smad4 indicates that all R-Smads and Smad4 are able to interact specifically with 5GC and SBE sites by means of a conserved binding mode, mostly using the β2-β3 hairpin. Only the long isoform of Smad2 shows additional contacts from residues in the E3 insert, exclusively present in this specific isoform ^22^.

The second hypothesis is related to DNA recognition. Although all Smad proteins can interact with SBE and 5GC motifs, the main difference observed in all MH1-DNA complexes is not in the recognition of different DNAs, but in the MH1 domain itself (monomer or dimer propensities). Whereas TGFβ-activated Smads and Smad4 interact with DNA as MH1 monomers, BMP-activated Smads form MH1 dimers by swapping the α1 helix between two monomers (Figure 4A). For a given Smad hetero-trimer, finding the optimal DNA sites genome wide must fulfill certain specific spatial requirements dictated by the MH1 structures (Figure 5D), which implies that not all theoretically available DNA motifs can be recognized. It also suggests that the presence of Smad4 in these complexes, always monomeric, provides some flexibility to the hetero-trimeric complex, even in the case of BMP-Smad4 complexes. This flexibility could explain why, for a given promoter, the distances between motifs are not strictly conserved among vertebrates. This versatility can also explain how so many different regions in the genome are bound by the different hetero-trimeric Smad complexes, as observed in ChIp-Seq data and also reported in the literature ^39,40^.

Third, the composition of Smad heterotrimeric complexes can be modulated by the association through dimers, and not only through MH2 domain interactions. It is well accepted that Smad associations are driven *via* direct contacts of the conserved MH2 domains of R-Smads and a single Smad4 protein, as shown in the crystal structures of various complexes of MH2 domains ^4,5,34,35^. R-Smad/Smad4 heterotrimers have been observed with over-expressed as well as with endogenous full-length proteins ^32,33^. However, despite the high level of MH2 domain conservation among different R-Smads, not all variants of Smad complexes seem possible or biologically supported, as for instance, a BMP-Smad/TGFβ-Smad/Smad4 complex or a complex made of two TGFβ-Smads and one BMP-Smad have never been experimentally detected in cellular experiments ^36-38^, suggesting that a second layer of selection might exist to favor the composition of some complexes over others. One of these selection rules could include holding one dimer of Smad1/5/8 MH1 domains and one monomer of either Smad2/3/4 (whose MH1 domains do not form dimers) thereby suggesting that these dimers may also occur in full-length Smad complexes, in native conditions. We propose that this characteristic has been among the keys to shaping two classes of R-Smad proteins since the origin of metazoans.

All findings available till now suggest that the selection of optimal DNA targets in a native context is the result of a collaborative approach between the different Smad complexes and bound cofactors. This selection process seems to be also modulated by the internal association of Smad proteins, where all components fit in order to fine-tune the context-dependent action of BMP and TGFβ signals. Certainly, additional experiments, as well as structures of full-length Smad complexes bound to DNA, will finally illustrate how these different layers of interactions are defined and modulated.

## Acknowledgments

We thank Dr. N. Berrow (IRB Barcelona, protein expression unit) for help with some DNA constructs and with protein purification. We also thank the EMBL staff for assistance at the HTX facility (Grenoble), the joint EMBL and ESRF group for access to synchrotron beamlines ID29, ID23-1 and ID23-2 and the staff at The Automated Crystallography Platform staff (IRB Barcelona-CSIC) and at the ALBA synchrotron (Barcelona) for access to the BL13-XALOC beamline. Thanks also go to Dr. J. Massagué for insightful suggestions and discussions, Drs. M. Díaz and M. Vilaseca (Mass Spectrometry Core Facility, IRB Barcelona) for support with the IM-MS data, Dr. M. Navia for suggestions on binding assays, and Dr. B. Brutscher, (Institut de Biologie Structurale, Biomolecular NMR Spectroscopy Group, Grenoble, France) for help with the acquisition of the NMR data at 800 MHz. We also thank J. Cordero for some preliminary experiments.

T.G. was a PhD student funded by a Severo Ochoa and by the BBVA. Z.K., R.F, R.P., and B.B were co-funded by the European Union’s Horizon 2020 research and innovation programme under the Marie Skłodowska-Curie COFUND actions of the EMBL, IRB Barcelona and the PROBIST and PREBIST Postdoc and Predoc Programmes (agreements EMBL_291772, IRBPostPro2.0_600404 and PROBIST_754510, and PREBIST_754558). M.J.M is an ICREA Programme Investigator. This work was supported by the Spanish MINECO program (BFU2014-53787-P and BFU2017-82675-P, M.J.M), IRB Barcelona and the BBVA Foundation. Access to the HTX facility at EMBL (Grenoble) was granted by the Horizon 2020 Programme i*NEXT* of the European Commission (grant 653706, title: Smad complexes), and to the NMR facility (Grenoble) by the Instruct Integrating Biology program (grant 2520, title: Monomer-dimer equilibrium in Smad proteins). Access to Bio-SAXS BM29 was part of the MX-1941 BAG proposal and to ALBA through the BAG proposal 2018092972.

We gratefully acknowledge institutional funding from the CERCA Programme of the Catalan Government and from the Spanish Ministry of Economy, Industry and Competitiveness (MINECO) through the Centres of Excellence Severo Ochoa award.

## Methods

Detailed methods on protein expression and purification, NMR experiments and analysis, crystallography experiments, TWIM-MS experiments, and SAXS data acquisition and analysis can be found in SI Appendix, Methods. The atomic coordinates have been deposited in the Protein Data Bank, Small-angle scattering data and models in the SASBDB database, and NMR assignments and chemical shifts in the Biological Magnetic Resonance Data Bank (BMRB).

## Supplemental Information

### Methods

#### Protein production and cloning

The Smad5 (Uniprot: Q99717-1, Ser9-Arg143), the Smad5_gly, Smad5_3 mutant construct and Smad8 (O15198-1, Thr14-Pro144) domains were cloned using synthesized DNA templates with optimized codons for Bacterial expression (Thermo Fisher Scientific) and confirmed by DNA sequencing (GATC Biotech). Proteins were expressed fused to an N-terminal His-tag followed by a TEV or 3C protease cleavage site in *E. coli* BL21(DE3) or C41(DE3) pLysS and purified following standard procedures ^13^. Unlabeled and labeled samples were prepared using Luria Broth (LB) and minimal media (M9) cultures, respectively (Melford). D_2_O (99.95%, Silantes), ^15^NH_4_Cl and/or D-[^13^C] glucose (Cambridge Isotope Laboratories, Inc) were used to prepare the labeled samples ^42^. Cells were cultured at 37°C to reach an OD_600_ range of 0.6-0.8. After induction with IPTG (final concentration of 0.4 mM) and overnight expression at 20°C, bacterial cultures were centrifuged and cells were lysed (EmulsiFlex-C3, Avestin) in the presence of Lysozyme and DNase I and in PBS buffer at pH 7.5. The soluble supernatants were purified by nickel affinity chromatography (HiTrap Chelating HP column, GE Healthcare Life Science) using an NGC™ Quest 10 Plus Chromatography System (Bio-Rad). Eluted proteins were digested with TEV or 3C proteases (at 4°C or room temperature respectively) and further purified by ion-exchange chromatography using a HiTrap SP HP and size-exclusion chromatography on a HiLoadTM Superdex 75 16/600 prep grade columns (GE Healthcare) equilibrated in 20 mM Tris-HCl buffer (pH 7.2), 80 mM NaCl and 2 mM TCEP.

Purified proteins were verified by Liquid chromatography-Mass Spectrometry (LC-MS) using an ACQUITY UPLC Binary Sol MGR LC system (Waters) equipped with a BioSuite Phenyl 1000Å column (Waters, 10 μm RPC 2.0×75 mm) at a flow rate of 100 μL/min. The column outlet was directly connected to the mass spectrometer, which acquired full MS scans (400-4000 m/z) working in positive polarity mode. Samples were eluted using a linear gradient from 2% to 5% B in 5 min and from 5% to 80% B in 60 min (A= 0.1% Formic Acid, FA, in water, B= 0.1% FA in CH3CN) and analyzed using the MassLynx™ Software, (V4.1.SCN704, Waters). The purity of the recombinant proteins was over 95%, as shown by the Mass Spec analysis.

#### Duplex DNAs

Duplex DNAs were annealed using complementary single-strand HPLC-purified DNAs. DNAs were mixed at equimolar concentrations (1 mM), heated at 90°C for 3 min and allowed to cool to room temperature for 2 h. DNAs (with and without fluorophores) were purchased at Biomers and/or at Metabion, Germany.

#### NMR chemical shift assignment and perturbation experiments

NMR data were recorded on a Bruker Avance III 600-MHz spectrometer equipped with a quadruple (^1^H, ^13^C, ^15^N, ^31^P) resonance cryogenic probe head and a z-pulse field gradient unit at 298 K. Backbone ^1^H, ^13^C and ^15^N resonance assignments were obtained by analyzing the 3D HNCACB and HN(CO)CACB experiment pair. Experiments were acquired as Band-Selective Excitation Short-Transient-type experiments (BEST) with TROSY and Non-Uniform Sampling (NUS) ^43,44^. This strategy allowed us to unambiguously assign 110 of the 121 possible amides (131 residues, 10 of them prolines). Comparison of the Smad5 MH1 (Cα and Cβ) chemical shifts to reference values, as well as the ^15^N edited-NOESY data, corroborated the presence of bound Zn^2+^ and of four helices and six strands, characteristic of the MH1 fold. The strands are ordered as three anti-parallel pairs: β1β_5_, β_2_β_3_, and β_4_β_6_. The presence of many long-range interactions confirmed that, in the absence of DNA, the structure of the MH1 domain is well defined in solution. Chemical shifts have been deposited in the BMRB (entry 27548). For the titration experiments, HSQCs were recorded using a Non-Uniform Sampling (NUS) acquisition strategy to reduce experimental time and increase resolution.

T1 and T2 relaxation measurements were acquired using standard pulse sequences ^43^. The rotational correlation time of the Smad5 MH1 domain (τc) was calculated assuming slow molecular motion, τc larger than 0.5 ns, and only J(0) and J(ωN) spectral density terms contributed to the overall value.

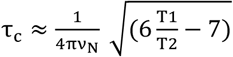 (6 - 7), where ν_N_ is the ^15^N resonance frequency.

Protein samples (400 μM for backbone experiments) were equilibrated in a 20-mM TRIS buffer containing 100 mM NaCl at pH7.2 supplemented with 10% D_2_O. Spectra were processed with NMRPipe ^45^ and MddNMR ^44^, and analysis was performed with CARA ^46^.

#### X-ray

High-throughput crystallization screening and optimization experiments were performed at the HTX facility of the EMBL Grenoble Outstation ^47^. Human Smad5 was concentrated to 5 mg/mL prior to the addition of the annealed DNAs (Metabion) dissolved in 20 mM Tris-HCl pH 7, 10 mM NaCl and 2 mM TCEP. The final molar protein DNA ratio was 1:1. Screenings and optimizations were prepared by mixing 100 nL of the complex solution and 100 nL reservoir solution in 96-well plates. Crystals of the complexes were grown by sitting-drop vapor diffusion at 4°C. Crystals were obtained with several DNAs of different lengths and in different conditions, and in all cases they were reproducible. Several datasets were acquired for the best diffracting crystals and were then analyzed. Final conditions for the three best diffracting complexes were optimized as follows:

Smad5: 0.2 M NaF, 0.1 M bis-tris propane pH 7.5, 20% PEG3350

Smad8: 0.2M NaF, 0.1 M bis-tris propane pH 8.5, 20% PEG3350

Smad5_3 chimera: 0.05 M sodium citrate pH 5.5, 22% PEG3350

Smad5_gly: 0.05 M Hepes pH 7.0, 21% PEG Smear Medium (PEG 2000, 3350, 4000, and5000 MME).

All crystals were cryoprotected in mother liquid supplemented with glycerol and frozen in liquid nitrogen. Diffraction data for Smad5 and Smad8 complexes were recorded at the ESRF in Grenoble (France) (beamline ID30a3) and Smad5_3 chimera and Smad5_gly data at the ALBA Synchrotron Light Facility (BL13-XALOC beamline), Barcelona, Spain. The data were processed, scaled and merged with autoPROC ^48^. Initial phases were obtained by molecular replacement using PHASER ^49,50^ from the CCP4 and PHENIX suites ^51,52^ (search model PDB code: 3KMP) with anisotropic correction. REFMAC ^53^ phenix.refine ^51^ and BUSTER ^54^ were employed for the refinement, and COOT ^55^ for the manual improvement of the models. For Smad5 mutants, the PDB-REDO server was used for the selection of data resolution cutoff (paired-refinement) and for the structure model optimization ^56^. Water molecules bound at the DNA-protein interface were selected when they participated in at least three hydrogen bonds (cutoff distance of 3.5 Å). Figures describing the structures were generated with UCSF Chimera^57^.

#### TWIM-MS experiments

Experiments were performed using a Synapt G1 HDMS mass spectrometer (Waters UK Ltd., Manchester, UK). Mass spectra were acquired by positive nano-electrospray ionization (ESI) using a Nanospray Triversa (Advion Biosciences Corpn., Ithaca, NY, USA) interface. To optimize the separation of the different conformers, traveling-wave drift times of selected ions corresponding to monomers and dimers of Smad MH1 domains (in 150 mM ammonium acetate buffer) were measured at wave heights of 7 V, 8 V, 9 V, and 10 V and at a velocity of 300 m/s. Data acquisition and processing were carried out using MassLynx (v4.1) software. Drift time calibration of the T-Wave cell was performed using β-lactoglobulin (monomer, 18 kDa, and dimer, 37 kDa) from bovine milk. Reduced cross-sections (Ω’) were calculated from published cross-sections ^58^ and subsequently plotted against final corrected drift times (tD). Calibration coefficients were determined applying an allometric y = AxB fit. Experimental cross-sections were determined by measuring the drift time centroid for the molecular-related ions by means of Gaussian fitting to the drift time distribution (Prism v6, GraphPad Software Inc., California, USA).

#### SAXS data

Data were collected on samples of Smad5 MH1 domain at protein concentrations ranging from 0.96 to 10 mg/mL. Only data from 0.96 to 3.41 mg/mL were used for the analysis. All samples were concentrated in 20 mM Tris buffer, 150 mM NaCl, and 2 mM TCEP, pH 7.2. Data were acquired at Beamline 29 (BM29) at the European Synchrotron Radiation Facility (ESRF; Grenoble, France). Protein samples were centrifuged for 10 min at 10,000*g* prior to data acquisition. Experiments on BM29 were collected at an energy of 12.5 keV and data were recorded on a Pilatus 1M detector at 10°C. For each sample and buffer, 10 exposure frames of 1 s were collected, and the exposure set was combined during data reduction to produce each SAXS curve. Buffer subtraction was performed after data reduction. Image conversion to the 1D profile, data reduction, scaling and buffer subtraction were done by the software pipeline available at the BM29 beamline. Further processing was done with the ATSAS software suite and Scatter ^59^. Guinier plot calculation (for the estimation of the radius of gyration Rg) was performed with PRIMUS, included in the ATSAS suite, using low q regions (qmax × Rg<1.3). Small-angle scattering data and models were deposited in the SASBDB database under the entry code SASDE32. Regarding the possible conformations in solution, the following three states were considered plausible: the crystallographic dimer; the closed monomer present in other crystal structures of MH1 domains; and an open monomer where the extended N-term helix was considered as flexible. Molecular modeling for the N-term helix structure was performed with the Rosetta modeling software suite, using the FloppyTail application ^60^ and starting from the MH1 crystal structures of Smad5 determined in this work (PDB: 6FZS). For closed monomer models, we used Modeller ^61^ and the Smad4 MH1 structure (PDB: 3QSV) as template, whereas the dimers were directly taken from the complexes of Smad5 (PDB: 6FZS). Trimers were generated using CNS. In all cases, DNA and water molecules were removed and secondary structure elements were restrained, except for the flexible N and C-terminal tails and the flexible N-term helix. For each state, one thousand conformers were simulated, in order to generate sufficient conformational sampling. Theoretical SAXS curves were calculated using CRYSOL ^62^ and compared to the experimental data, selecting the linear combination of the three states with the lowest chi-squared, as implemented in ATSAS and in in-house scripts.

The chi-squared metric for N data points was calculated using the equation:

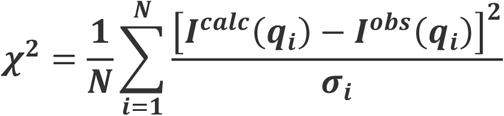

#### ChIP-Seq statistical analysis

Chip-Seq datasets (SRP179614 for Smad3 and GSM1810980 for Smad1/5) were downloaded from the NCBI SRA ^63^ and GEO ^64^ data repositories.

The Smad3 data was downloaded, the fastq format data was extracted with *fastq-dump* (**SRA Toolkit**, SRA toolkit, SRa Toolkit Development Team, http://ncbi.github.io/sra-tools/). Bowtie2 ^65^ was used to align the fastq reads against the mouse mm10 assembly, and sambamba ^66^ was used to sort and remove duplicate and unmapped reads. Peak calling was performed with macs1.4, for consistence with the Smad1/5 dataset ^67^.

The Smad1/5 data was downloaded from the GEO database. As the original data was aligned against the mm9 genome assembly, the UCSC liftOver software ^68^ was used to convert the coordinates to the mm10 assembly.

All ChIP-seq peaks were normalized to 200bp centered with respect to the peak center, using a custom python script and all regions were scanned for SBE+5GC motifs (SBE: GTCTG, 5GC: GGCTG, GGCGC and GGCCG) and the number of motifs per Kb as well as the distance of each motif to the nearest one in the same band downstream the genome were determined. By doing this we obtained a number of distances per band which is equal to the number of detected motifs minus 1.

The Statistical analysis was performed using R language, version 3.6.3 (R Core Team (2017) R: A Language and Environment for Statistical Computing). The statistical analysis used the R built-in functions *ad*.*test* for the Anderson-Darling normality test and *wilcox*.*test* for the Wilcoxon rank sum test. Plots display as Figure 5A and 5B were generated with the ggplot2 package.

For promoter analysis, we assigned each ChIP-Seq peak to the nearest annotated Transcription Start Site, and then extracted the ones related to the genes of interest. The gene names and the coordinates for the bands are included in Supplementary Table S5. The Smad3 ChIP-seq was used to extract bands for the TGF-β regulated genes and the Smad1/5 ChIP-Seq was used for the BMP regulated ones.

## Supplementary Figures

**Supplementary Figure S1.**
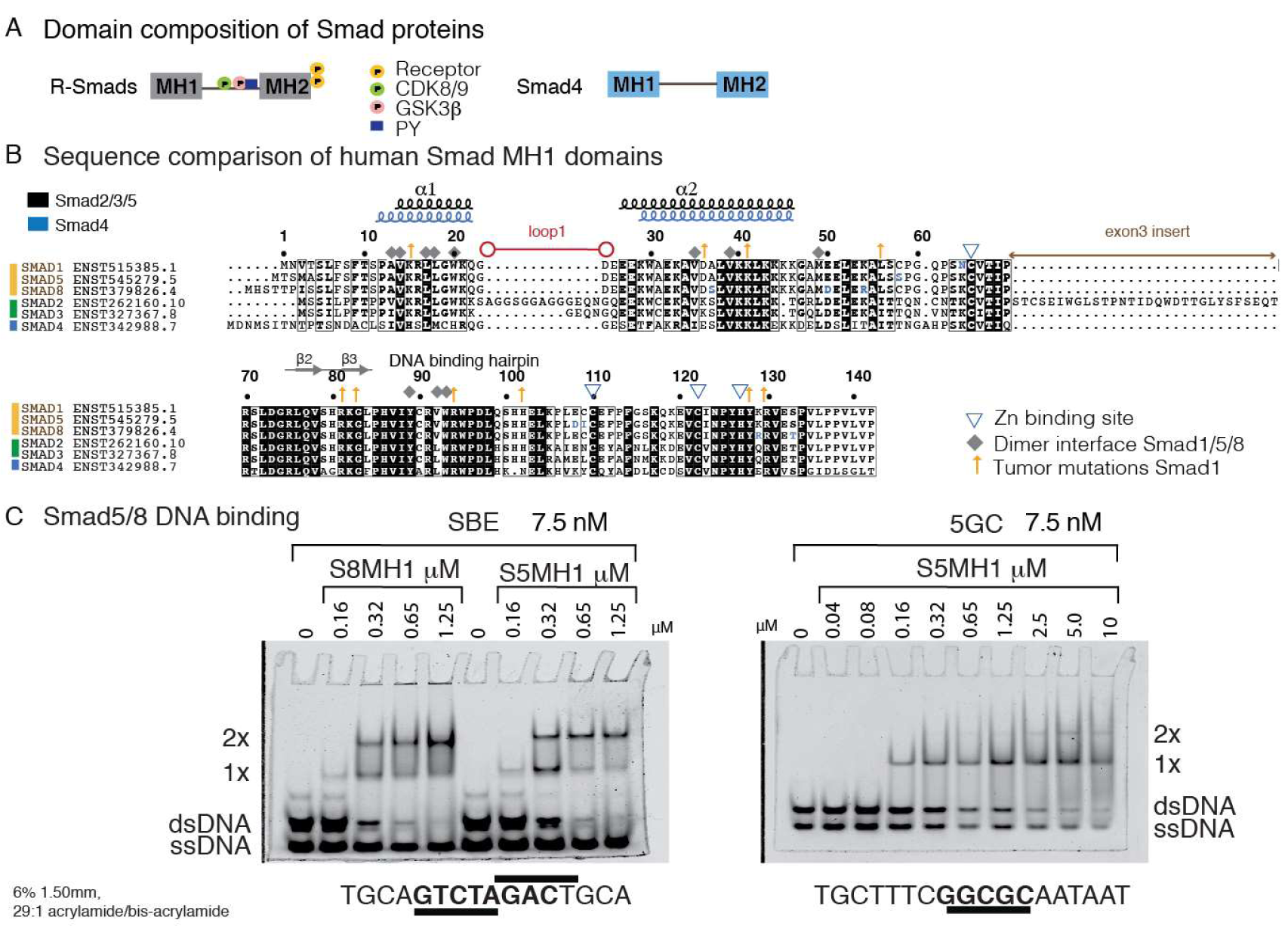
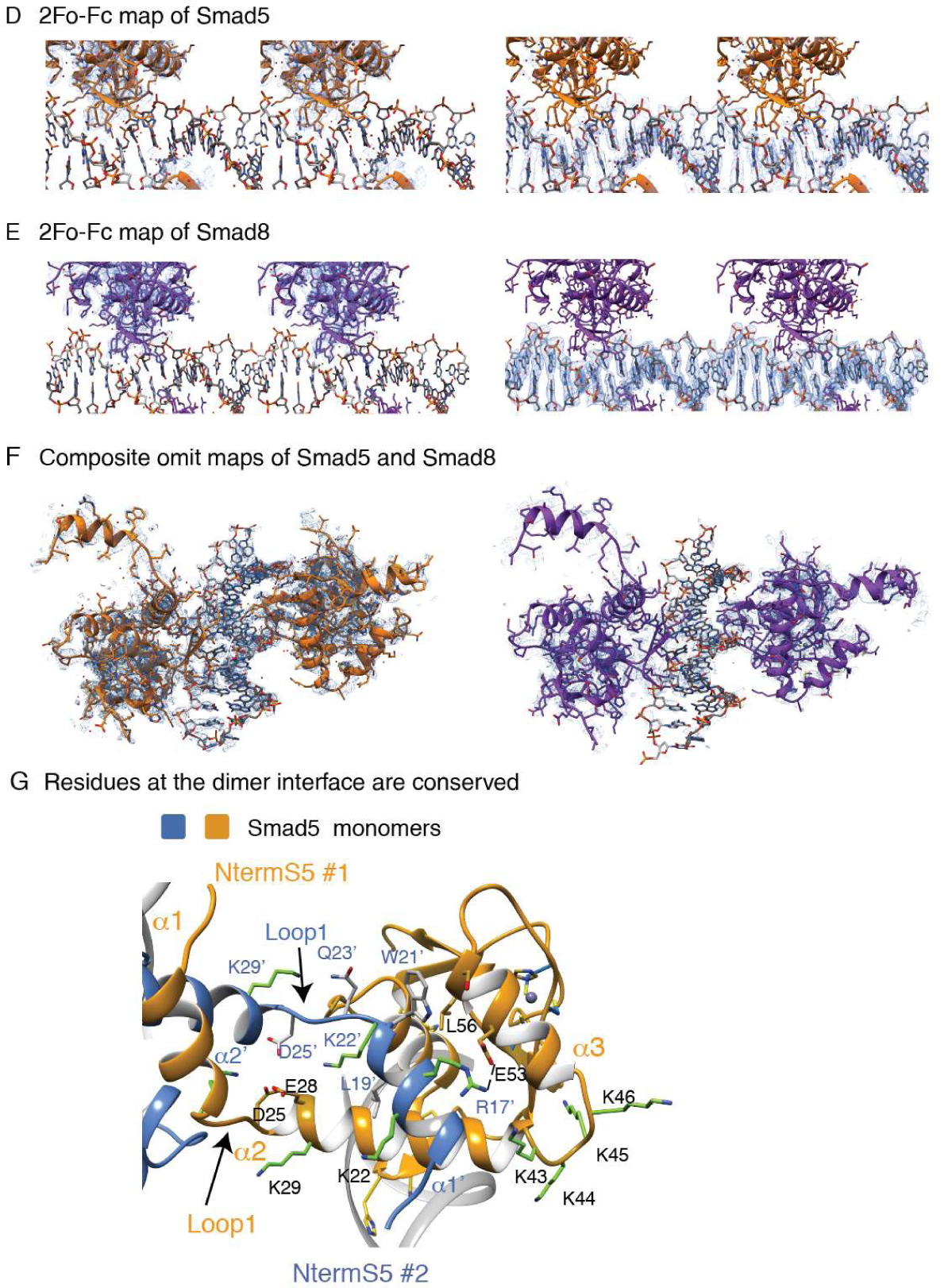
A. Domain composition of R-Smads and Smad4. Different phosphorylation sites of R-Smads are indicated as yellow (receptor), green (CDK8/9) and pink (GSK3) circles. B. Sequence alignment of the R-Smads and the Co-Smad Smad4 MH1 domain (human proteins), where strictly conserved residues are highlighted. The four residues that coordinate Zn are indicated with a triangle. The residues that define the first two helices (as observed in the structures of R-Smads and in Smad4) and the DNA binding hairpin are represented on the top of the alignment. The variable length of loop1 is indicated with a red line. Residues participating in the dimerization interface of BMP-activated Smads are indicated as gray diamonds. Tumor mutations described for Smad1 are indicated by arrows. The insert corresponding to the Smad2 long isoform is labeled. Alignments were generated with MAFFT ^69^ and BoxShade server. C. EMSA assays showing the interactions of Smad5 and Smad8 with the SBE sequence and the Smad5 interaction with the 5GC sequence. The binding affinity is similar for both sequences. DNA sequences are shown at the bottom, DNAs were at a 7.5nM fixed concentration and MH1 domain concentration for each lane is shown on top. D. Section of the electron‐density map (stereo-view) contoured at 1.0σ for the Smad5 MH1 complex. The map was calculated with coefficients 2Fo – Fc, and the refined model (ribbon) is superimposed. The density map is shown for the protein or for the DNA. Some residues and elements of secondary structure are indicated. E. Section of the electron-density map (stereo-view) representing the protein-DNA complex of Smad8 as in panel A. F. Composite OMIT maps for the Smad5 MH1 domain (left) and the Smad8 MH1 domain (right). G. Smad5 residues located at the dimer interface are labeled in blue and black. These residues are also conserved in Smad1 and Smad8.

**Supplementary Figure S2.**
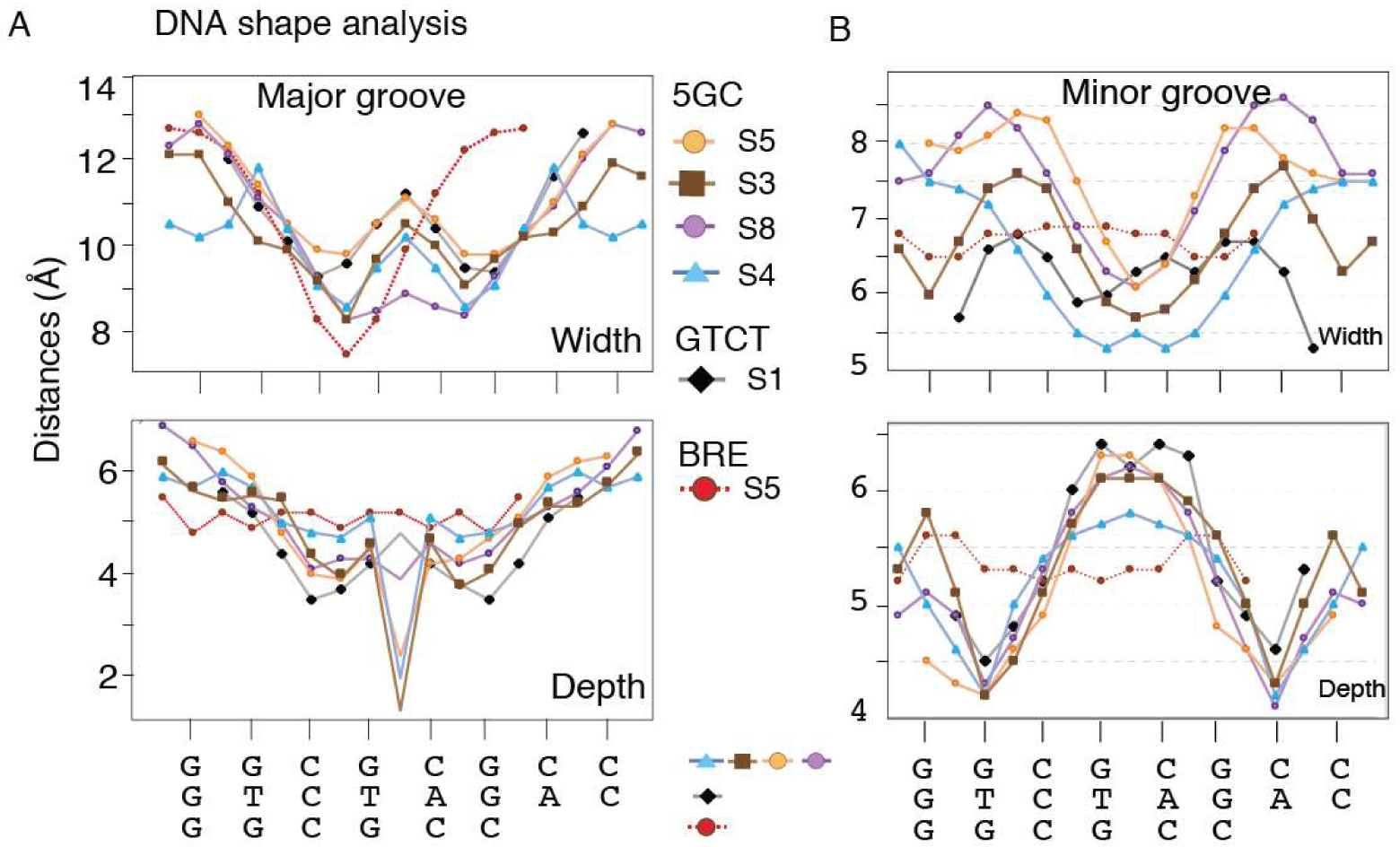
A. Major groove width and depth analyses of the DNA bound to Smad5 (orange), Smad8 (lavender), Smad3 (brown), Smad4 (blue), Smad1 (black) and Smad5-BRE complex (red). The values were calculated using Curves+ ^19^. B. Minor groove width and depth analyses of the DNA bound to Smad5 (orange), Smad8 (lavender), Smad3 (brown), Smad4 (blue), Smad1 (black) and Smad5-BRE complex (red). The values were calculated using Curves+ ^19^.

**Supplementary Figure S3.**
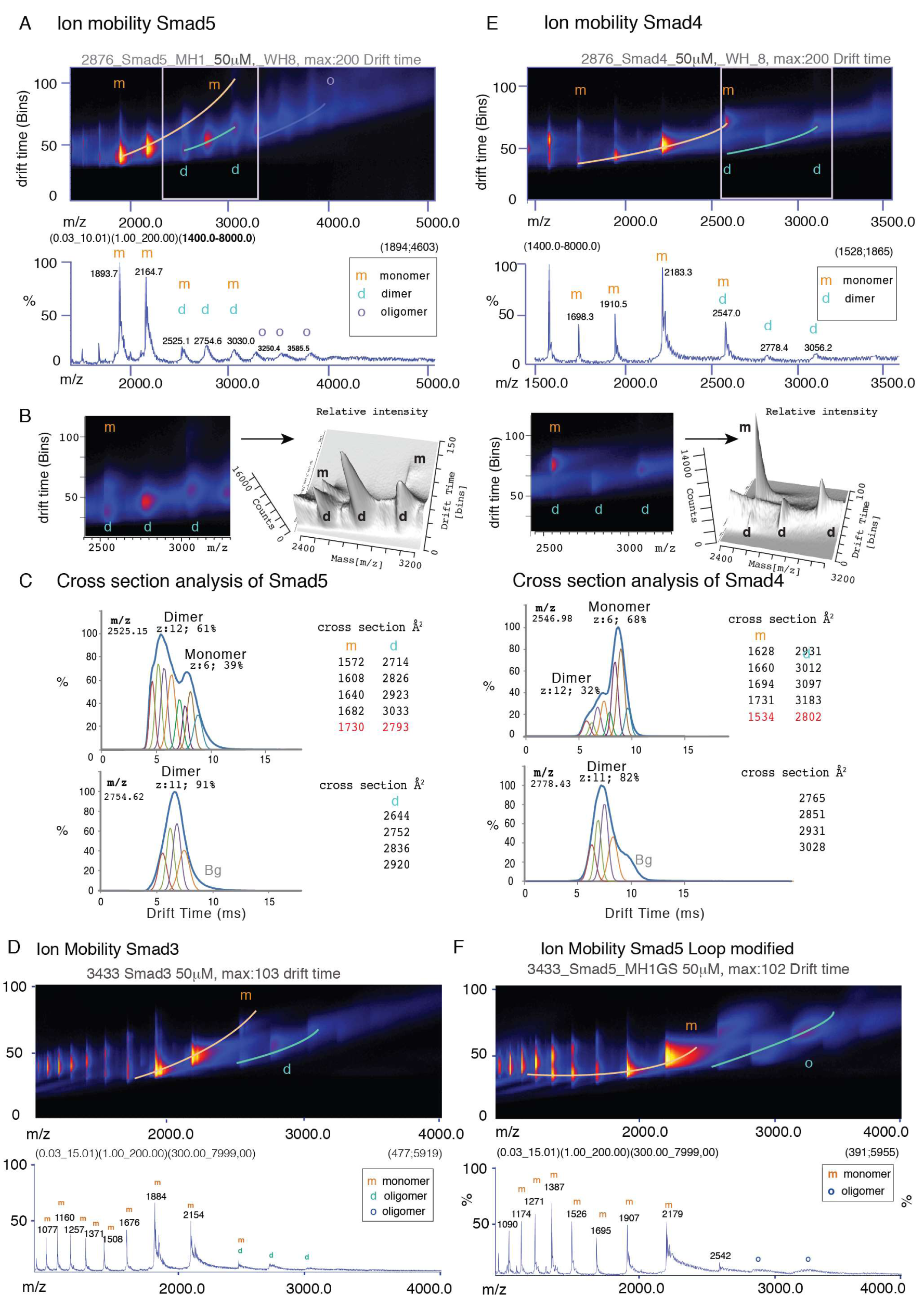
Multidimensional IM-MS data of Smad5, Smad 3 and Smad4 MH1 domains. A. Plot of the mobility drift time versus *m/z* for the Smad5 MH1 domain (aqueous solution, 200 mM ammonium acetate buffer) showing the multidimensional data, (instrumental settings are provided in table S3). The mass spectrum from all ions shown below. Peaks corresponding to monomers (m) and dimers (d) present in the gas phase are labeled. B. Close-up representation (left: 2D contour plot and right, 3D plot) of the conformations corresponding to monomers and dimers that are present at a given m/z value but that are separated by ion mobility. C. Ion mobility time distributions for three selected charge states. Collision cross-sections and the conformations used for the fitting. Calibration curves using standard proteins were used prior to the cross-section analysis. D. Equivalent results to panel A are shown for Smad3 E. Equivalent results to panels A, B and C are shown for Smad4 F. Equivalent results to panel A are shown for Smad5_gly mutant with the modified loop1.

**Supplementary Figure S4:**
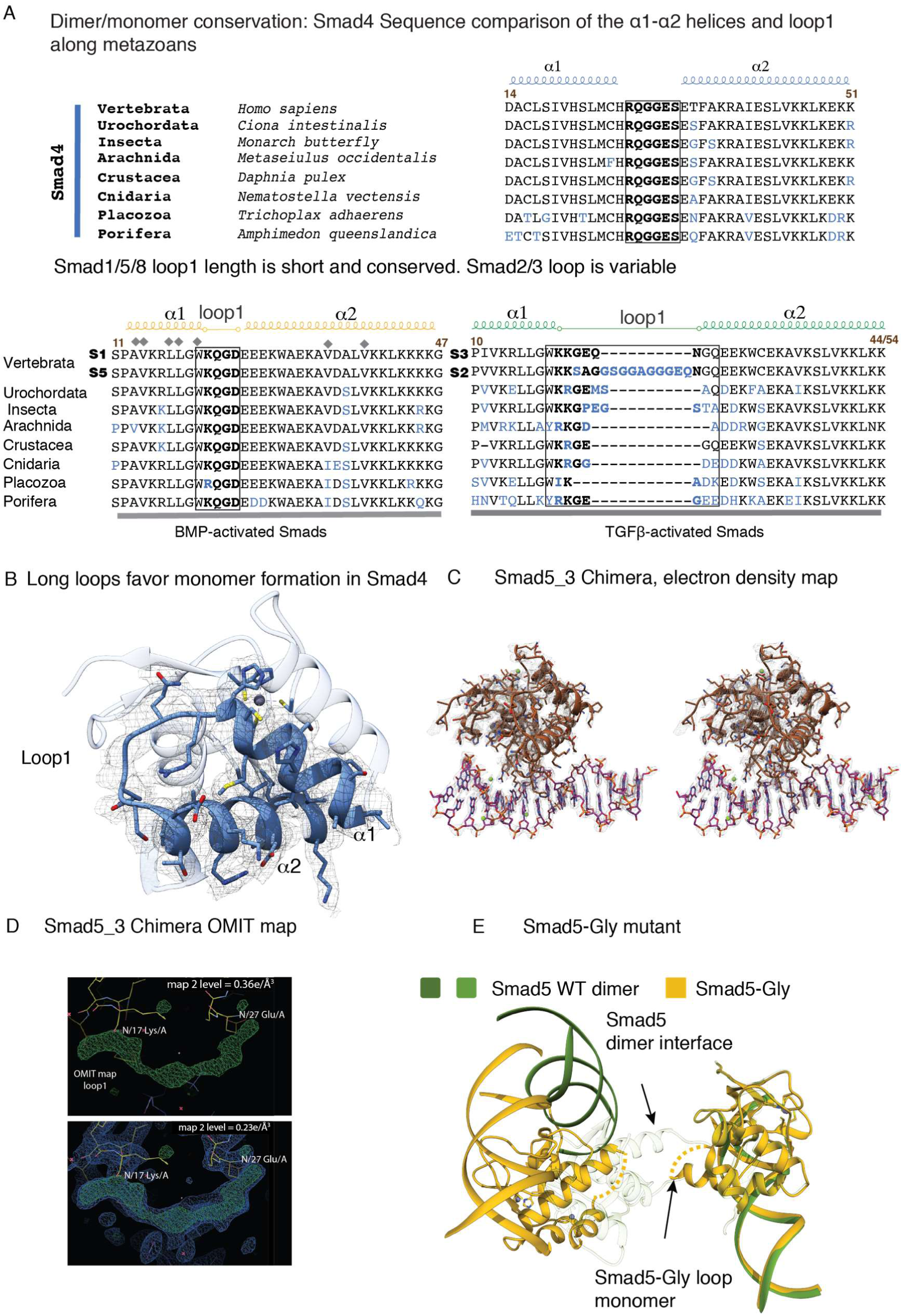
Sequence conservation and dimer interface. A. Comparison of the а1-а2 regions along different metazoans. The specific organisms used in the comparison are indicated in the Smad4 alignment. B. Smad4 MH1 monomer (pdb: 5MEY). Smad4 does not show side-chain contacts stabilizing the loop1 orientation, in opposition to Smad3, while still maintaining the monomeric arrangement. Some elements of the secondary structure are labeled. C. Section of the electron‐density map (stereo-view) contoured at 1.0σ for the Smad5_3 chimera complex. The map was calculated with coefficients 2Fo – Fc, and the refined model (ribbon) is superimposed. D. OMIT and electron‐density maps corresponding to loop1 (Smad5_3 chimera). The refined model is superimposed and some residues are labeled. E. Superposition of the Smad5 dimer and the Smad5_gly monomer with Loops 1 labeled. The loop in the Smad5_gly monomer is indicated as a dotted line since its corresponding electron‐density is not well defined.

## Supplementary Tables

**Supplementary Table S1.** Analysis of Water Molecules (WM) that mediate interactions between MH1 domains and the GGCGC DNA. Color code: <u>**Underlined**</u>: WM between Monomer B and DNA **Black:** WM between Monomer A and DNA **Gray:** Crystallographic WM between Monomer A and DNA

| Protein complex | Water number | Atoms involved, chain identifier, BB: backbone distances |  |  |  |
| --- | --- | --- | --- | --- | --- |
| Smad5 | **W23** | PO<sub>4</sub> of G8 (C)<br>2.63 Å | Gln77 (A)<br>2.78 Å | BB His80, (A)<br>3.1 Å | BB Ser79 (A)<br>3.5 Å |
|  | **W24** | PO<sub>4</sub>, of G8 (C)<br>2.7, 3.4 Å | BB Gln77 (A)<br>2.7 Å |  |  |
|  | **W36** | PO<sub>4</sub> of G10 (C)<br>2.7 Å | BB Asp73 (A)<br>3.0 Å | BB Leu72 (A)<br>3.5 Å |  |
|  | **W53** | PO<sub>4</sub>, G8 (D)<br>2.8, 3.6 Å | BB Gln77 (B)<br>2.8 Å |  |  |
|  | **W58** | PO<sub>4</sub> of G8 (D)<br>2.7 Å | Gln77 (B)<br>2.9 Å | BB His80 (B)<br>3.4 Å | BB Ser79 (B)<br>3.3 Å |
|  | **W100** | T1' Carbonyl<br>3.5 Å | Glu51 (A) 3.0 Å | BB Gly47 (A)<br>3.2 Å |  |
|  | **W118** | PO<sub>4</sub> of G8 (D)<br>2.76, 3.40 Å | BB Ser79 (B)<br>3.5 Å |  |  |
|  | **W122** | PO<sub>4</sub> of C9 (D)<br>2.7 Å | BB Arg75 (B)<br>3.0 Å |  |  |
|  | **W130** | N7 of G8 (D)<br>2.8 Å | His80 (A)<br>3.6/3.3 Å | W85<br>3.1 Å | W59<br>3.1 Å |
|  | **W131** | N7 of G6 (C)<br>3.3 Å | N4 of C7 (C)<br>2.9 Å | Gln77 (B)<br>3.1 Å | W132,<br>3.3 Å |
| Smad8 | **W49** | PO<sub>4</sub> of G10 (C)<br>3.2 Å | Asp73 (A)<br>2.8 Å | BB Asp73 (A)<br>3.6 Å | Ser71 (A) 3.5 Å |
|  | **W90** | PO<sub>4</sub> of C9 (D)<br>3.0 Å | Lys41 (B)<br>2.8 Å | W69<br>3.2 Å |  |
|  | **W91** | G8 PO<sub>4</sub> (D)<br>2.7 3.33 Å | Ser37 (B)<br>3.3 Å | W69<br>3.1 Å |  |
|  | **W95** | PO<sub>4</sub> of C9 (D)<br>3.1 Å | BB Pro69 (A)<br>3.6 Å |  |  |
|  | **W96** | PO<sub>4</sub> of G10 (D)<br>2.8 Å | BB Leu72, (B)<br>3.5 Å | BB and Asp73 (B)<br>3.1 Å, 3.6 Å | W108,<br>2.8 Å |
|  | **W97** | PO<sub>4</sub> of C9 (D)<br>2.8 Å | BB Arg75 (B)<br>3.1 Å |  |  |

**Supplementary Table S2.** RMSD values obtained from the comparison of several MH1 domain. RMSD values for the Smad5 MH1 domain superimposed to Smad8, Smad1 and Smad4 MH1 domains. Compared residues used for the fit are indicated in parenthesis. The PDB IDs are indicated for Smad1 and Smad4.

| **RMSD - Cα** | **Smad8**<br>**(5-135)** | **Smad1:3KMP**<br>**(9-132)** | **Smad4:3QSV**<br>**(8-135)** |
| --- | --- | --- | --- |
| **Smad5**<br>**(10-132)** | 0.25 Å | 0.30 Å | 0.60 Å |

**Supplementary Table S3.**
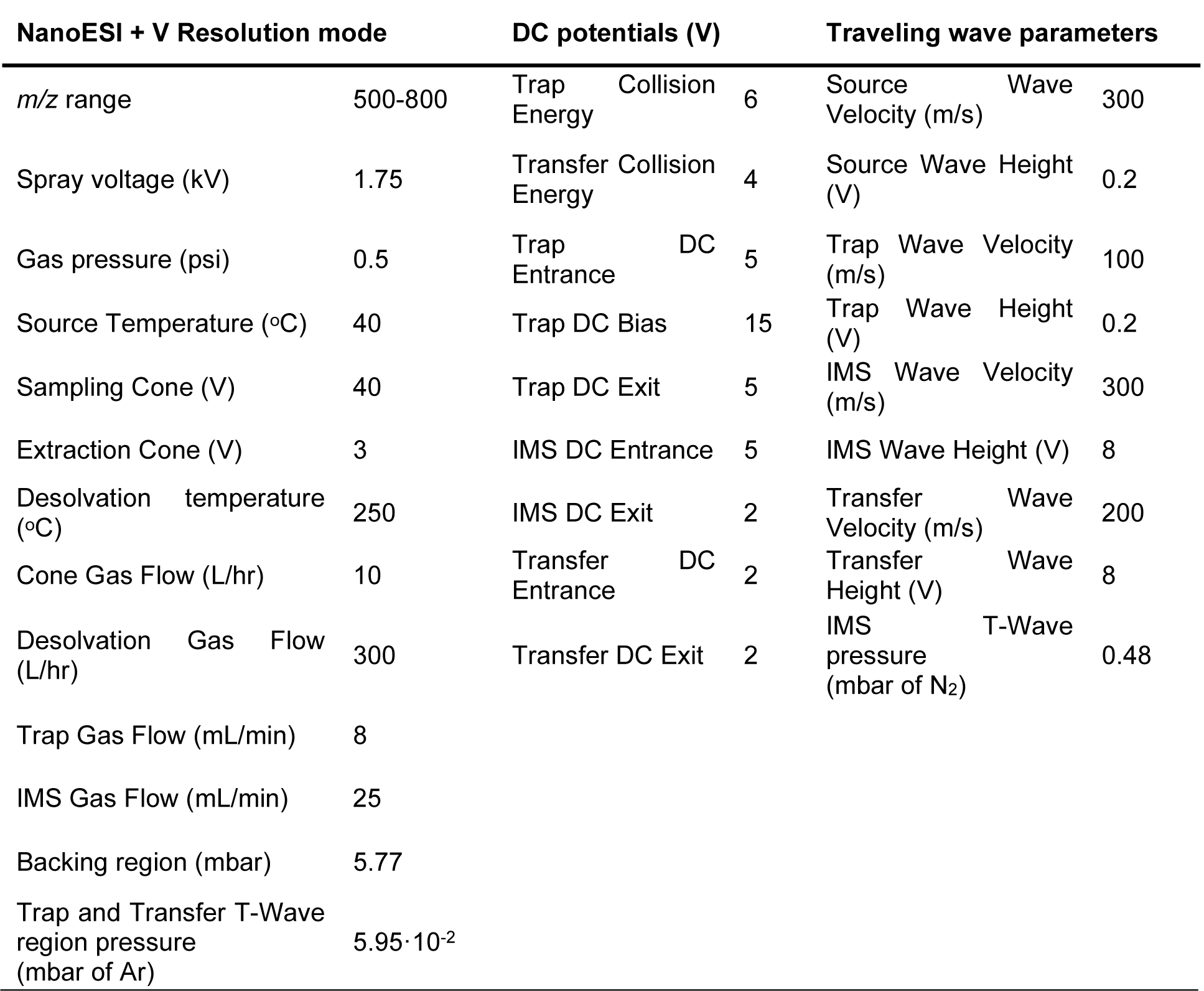
Experimental parameters used for ion mobility. Experiments acquired on a SYNAPT G1-HDMS (Waters, Manchester, UK)

**Supplementary Table S4.**
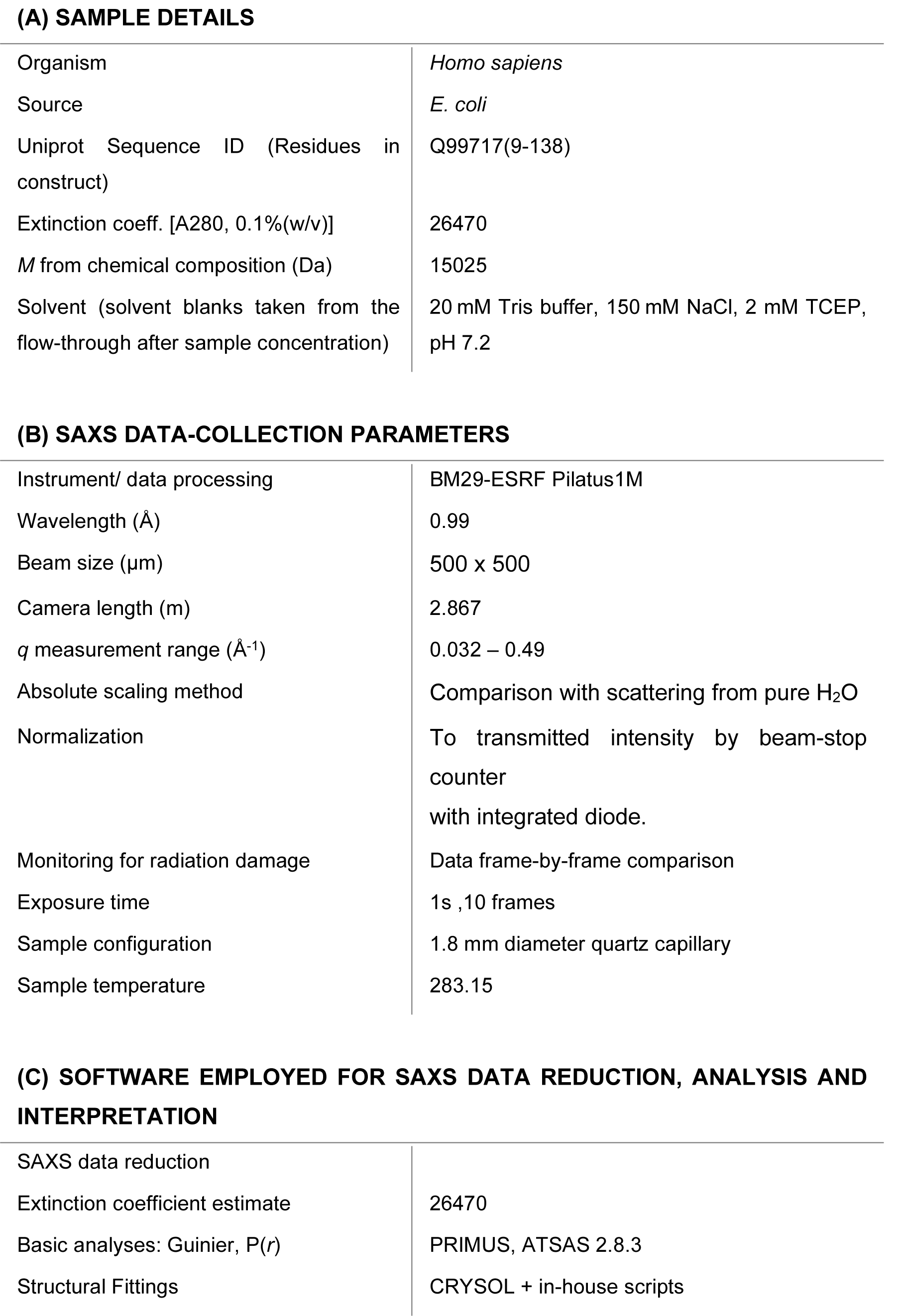
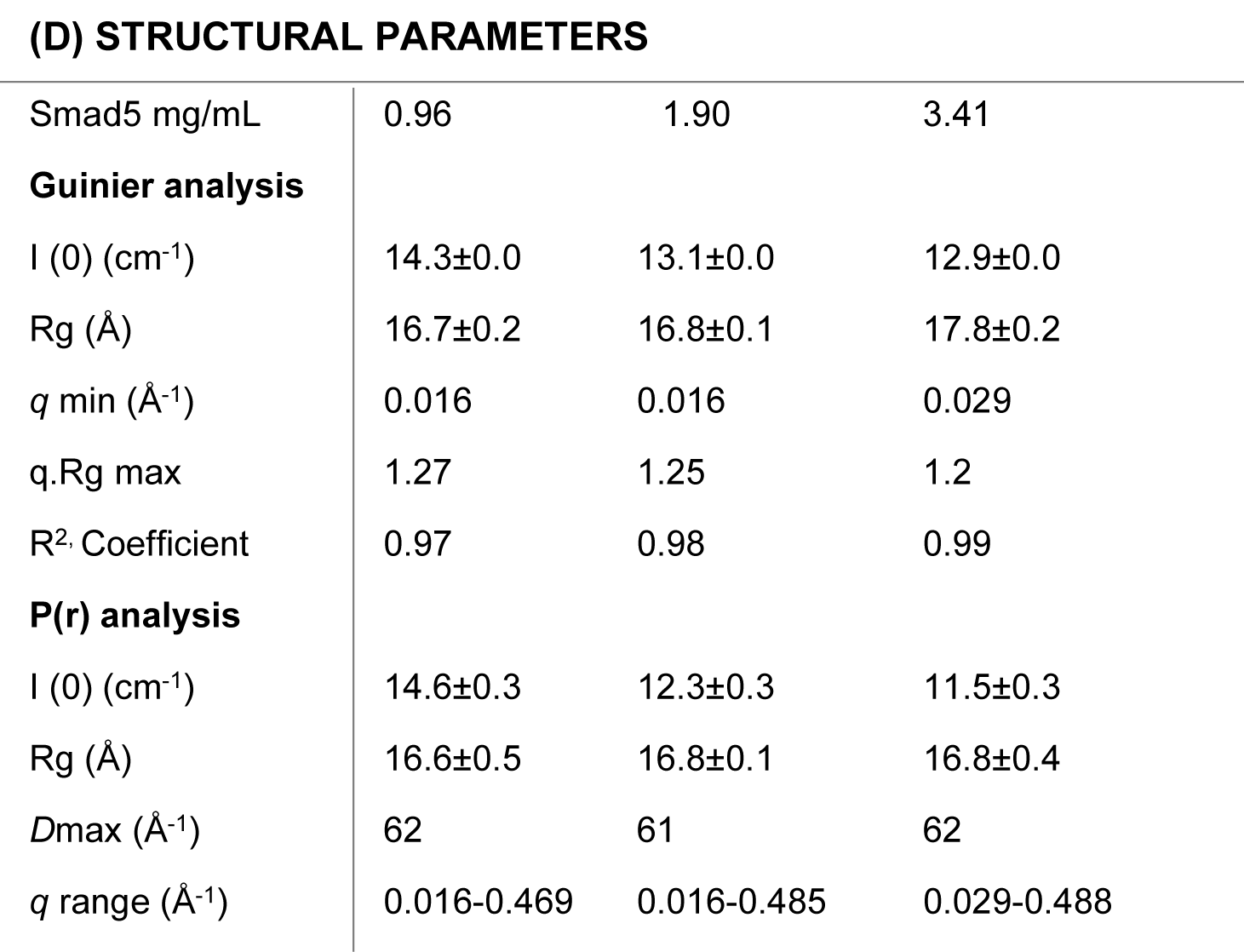
SAXS data and acquisition parameters for Smad5.

**Supplementary Table S5.**
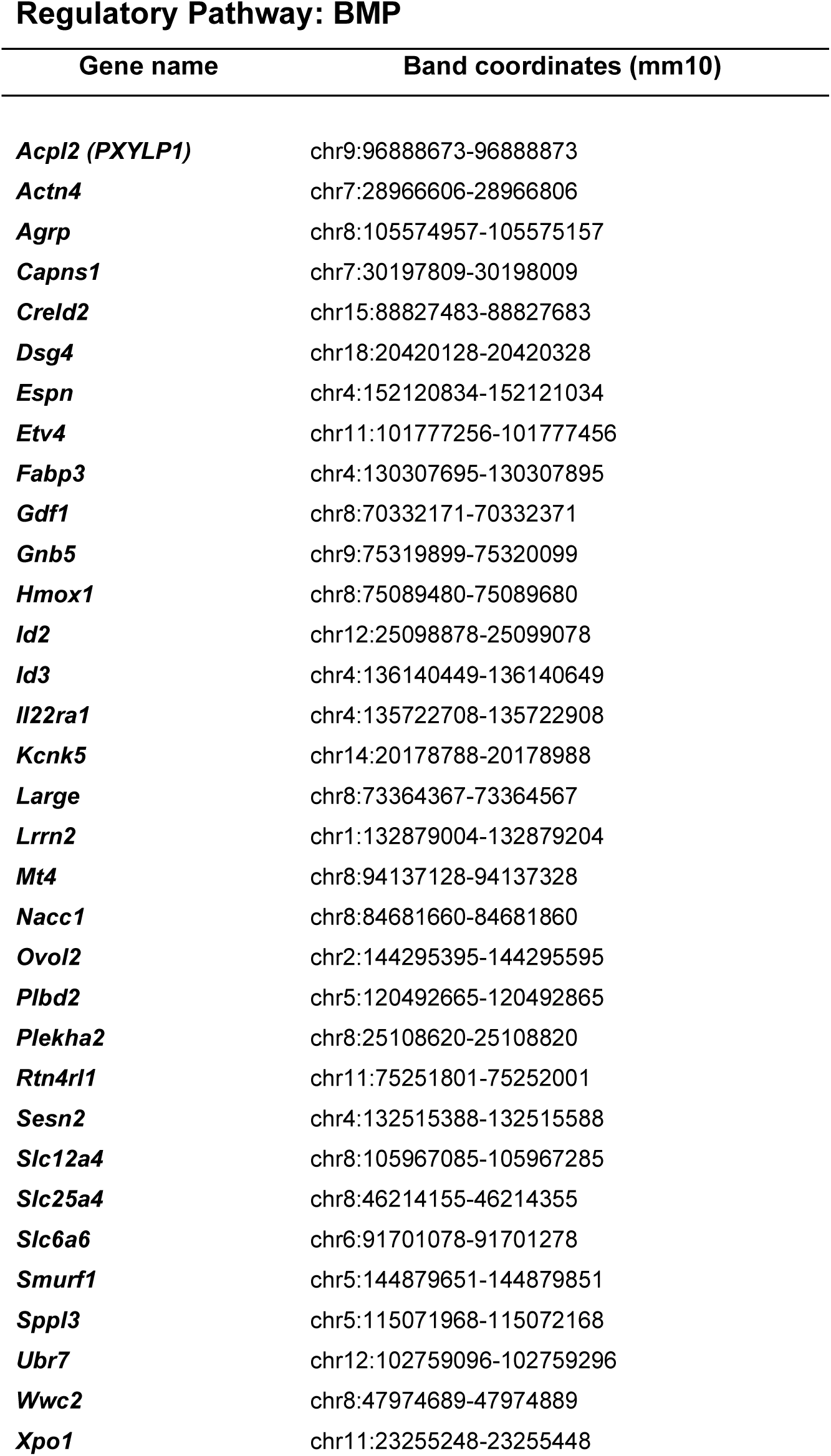
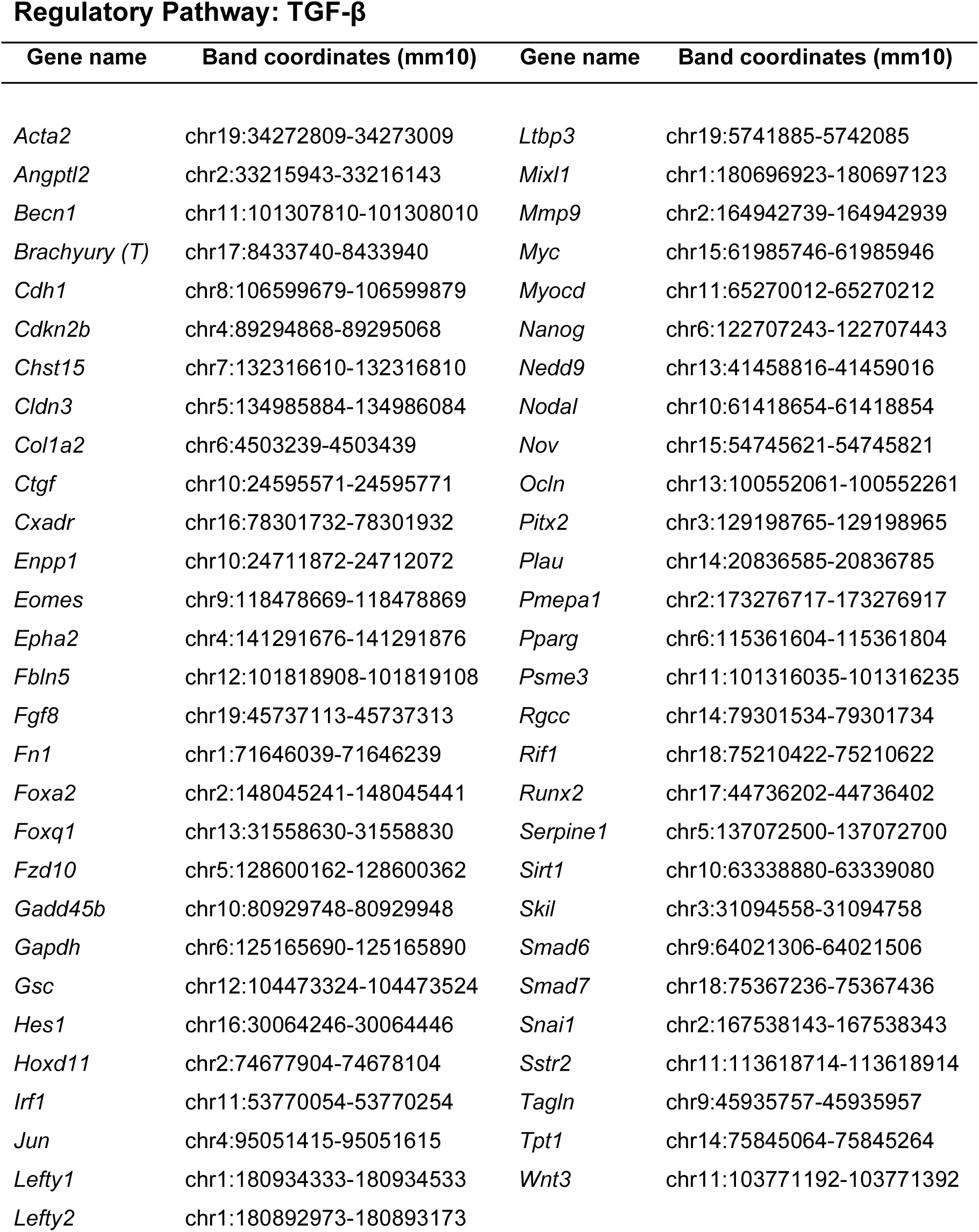
ChIPseq bands used for promoter analysis Regulatory Pathway: BMP.

## Footnotes

Footnotes Supplemental information is available for this article.

## Notes

### Competing Interest Statement

The authors have declared no competing interest.

### Summary of Updates

Modified the SAXS data table to follow the recent recommendations, removed the thermal unfolding experiments and included the Bioinformatic analyses as suggested by the reviewers. The latter includes the analysis of motif distribution TGF-B and BMP regulated genes in datasets available in the literature. Minor modifications were included in some figures as recommended.

